# PAbFold: Linear Antibody Epitope Prediction using AlphaFold2

**DOI:** 10.1101/2024.04.19.590298

**Authors:** Jacob DeRoo, James S. Terry, Ning Zhao, Timothy J. Stasevich, Christopher D. Snow, Brian J. Geiss

## Abstract

Defining the binding epitopes of antibodies is essential for understanding how they bind to their antigens and perform their molecular functions. However, while determining linear epitopes of monoclonal antibodies can be accomplished utilizing well-established empirical procedures, these approaches are generally labor- and time-intensive and costly. To take advantage of the recent advances in protein structure prediction algorithms available to the scientific community, we developed a calculation pipeline based on the localColabFold implementation of AlphaFold2 that can predict linear antibody epitopes by predicting the structure of the complex between antibody heavy and light chains and target peptide sequences derived from antigens. We found that this AlphaFold2 pipeline, which we call PAbFold, was able to accurately flag known epitope sequences for several well-known antibody targets (HA / Myc) when the target sequence was broken into small overlapping linear peptides and antibody complementarity determining regions (CDRs) were grafted onto several different antibody framework regions in the single-chain antibody fragment (scFv) format. To determine if this pipeline was able to identify the epitope of a novel antibody with no structural information publicly available, we determined the epitope of a novel anti-SARS-CoV-2 nucleocapsid targeted antibody using our method and then experimentally validated our computational results using peptide competition ELISA assays. These results indicate that the AlphaFold2-based PAbFold pipeline we developed is capable of accurately identifying linear antibody epitopes in a short time using just antibody and target protein sequences. This emergent capability of the method is sensitive to methodological details such as peptide length, AlphaFold2 neural network versions, and multiple-sequence alignment database. PAbFold is available at https://github.com/jbderoo/PAbFold.

## Introduction

Understanding where and how an antibody binds to its target protein is important for understanding how the antibody performs its function, whether that function is neutralizing a pathogen during an immune response, binding an epitope in immunoassays, or labeling a target molecule in a live-cell imaging experiment. However, determining the binding epitope of an antibody can be a time and labor-intensive endeavor with significant expense. Traditionally, antibody epitopes on target proteins have been identified by performing deletion analysis on the target protein to determine if the antibody loses reactivity for the deletion mutants in various immunoassays, which provides the general region of the target protein the antibody binds to. With the advent of widely available chemical peptide synthesis, sequence-specific synthetic peptides can be used for competitive immunoassays (such as enzyme-linked immunosorbent assays (ELISA)) to establish sequences that can effectively compete with the antigen for antibody binding. Peptide mapping experiments are a powerful method for determining the fine sequence of linear antibody epitopes, but these experiments can be relatively expensive and the time between experimental design and data acquisition can be weeks to months due to the need to design and chemically synthesize peptides. Once a peptide has been identified that binds with high affinity and specificity to an antibody antigen binding fragment (Fab), crystal structures can be determined that demonstrate intermolecular interactions between the peptide and antibody. These can then provide a molecular-level explanation for an antibody’s binding mode. Finally, with the advent of rapid single B-cell sequencing technologies to analyze humoral immune responses towards vaccination or infection, determining where specific antibody clones bind on an antigen becomes even more challenging due to the need to isolate or synthesize specific antibody genes, produce antibodies, and then perform deletion or epitope mapping experiments described above to fully understand how and where antibodies bind. These challenges make determining antibody epitopes expensive and time-consuming and limit the number of antibodies that are characterized in detail.

Antibodies that bind to linear epitopes represent an important subset to molecular biology, as they can be added to recombinant proteins for use in various types of immunoassays. By definition, a linear epitope is a binding site on an antigen that is recognized by the primary structure or contiguous linear sequence of amino acids. A number of linear epitope specific antibodies have been developed for use in various immunoassays (ELISA, western blot, immunofluorescence, etc.). The development of computational methods for linear epitope determination could increase the number and quality of new linear epitopes available to the field. Most epitope prediction tools (such as BepiPred (1), ElliPro (2), and ABCpred (3)) are generally designed to predict regions of an antigen that could be recognized by any antibody rather than a specific antibody. These programs also provide no insight into the structural match of the epitope and antibody, potentially making decisions without key structural information that otherwise may be relevant. The challenge in predicting epitopes for a *specific* antibody lies in the complexity of protein-protein interaction dynamics, which includes conformational changes, binding affinities, and thermodynamic stability. Structure based approaches including HADDOCK (4, 5) and ZDOCK (4, 6) can be used to dock peptides into antibody structures, but these require known peptides for binding. Significant progress has been made to address this problem via deep learning: some of the new and exciting tools are GearBind (7), PALM and A2binder (8), and DSMBind (9). We point the reader to this review for an excellent overview of some of the tools that have existed for some time, along with a comparison of these tools (10).

Determining antibody-epitope interactions is, at its most basic level, a structural biology problem. Determining what molecular interactions are present between an antibody and its antigen can define the epitope, determine what portions of the epitope and CDR sequences are responsible for molecular interactions, and provide clues to antibody specificity and affinity. With the advent of highly accurate structural predictions, including the AlphaFold2 (AF2) neural networks (11, 12), the ability to accurately predict protein structures, and potential protein-protein interactions, has dramatically increased. AlphaFold2 was trained on existing protein structures and can effectively model new protein structures.

Numerous antibodies, antibody Fab regions, and other related constructs with bound target peptides or proteins have been crystalized and deposited into the Protein Data Bank (PDB) (for example (13–16)). These PDB entries represent a valuable training set that may increase the likelihood that AlphaFold2 can successfully predict the structure for antibody-epitope complexes (12, 17–19). The authors of AlphaFold2 multimer (12) comment on the difficulty of predicting antibody-epitope complexes, and results for this are indeed mixed at best (17–19). One way in which this current report is distinct is our focus on linear epitopes. We hypothesize that the lack of strong competing structure within the short peptide may boost AF2 prediction of scFv-epitope binding predictions relative to conformational epitopes. This problem has precedent, as AlphaFold2 has previously been used to study the interactions between proteins and peptides (17, 18). AlphaFold2’s ability to correctly dock independent protein chains can be repurposed to predict how strongly two proteins interact together and extends to predicting the interaction between an antibody and short flexible peptides (linear epitopes) drawn from a larger protein antigen.

To maximize compute efficiency, it is helpful to minimize the size of the system subject to structure prediction. The computational expense of AlphaFold2 scales with the square of the length of the concatenated sequences involved. Fortunately, with respect to epitope specificity, antibody constant domains are less critical than the CDR loops and the remainder of the variable domain framework regions. Antigen binding by antibodies is primarily dictated by the antigen binding fragment (Fab) containing the variable light (VL) and variable heavy (VH) fragments. Conversion of full antibody sequences into single chain variable fragments (scFv) can significantly reduce structure prediction complexity and compute time. A wildtype scFv sequence can easily be generated directly from translated antibody heavy and light chain DNA sequences. Briefly, the sequences are first divided into framework and complementarity determining regions (CDRs) using Kabat (20) or IMGT (21) nomenclature. A flexible linker sequence (GGGGSGGGGSGGGGS, 15 a.a.) is then added between the new C-terminus of the truncated light chain and the original N-terminus of the shortened heavy chain to generate a single protein sequence that incorporates both antigen-binding chains. The resulting fusion protein often functions in a similar fashion to the original antibody. Another well-known protein engineering strategy for antibodies is “loop grafting”, where the CDR loops from one antibody are grafted onto a different framework region. We have recently used this approach to develop scFvs with improved *in vivo* performance (22). The structures of the novel scFv chimeras can be rapidly and confidently predicted by AlphaFold2 due to their small size and the extensive immunoglobin representation within sequence databases and the PDB. Excluding the time needed to obtain a multiple sequence alignment (MSA), predicting the structure for a single scFv in complex with a 10-a.a. peptide requires only 1.5 minutes on an NVIDIA A5000 graphics processing unit (GPU). This modest compute time allows a GPU-laden server or workstation to handle large-scale structure prediction of hundreds of related systems. As for the MSA input, a high quality MSA can quickly be obtained via ColabFold (23), which relies on the MMseqs2 MSA server. In our workflow, we repeatedly predict the structure for a fixed single scFv sequence in complex with varying peptide partners. In this case, we do not expect the peptide portion of the MSA to be useful. Therefore, to avoid sending hundreds of nearly identical MSA requests to MMseqs2 MSA server, and to avoid varying information in the MSA, we slightly modified the LocalColabFold code to include the option to cache the MSA (install available on the GitHub). We generate one cached MSA per epitope scan, where each residue in the query peptide is a glycine.

Several recent papers have attempted to use AlphaFold2 to identify antibody epitopes (24–26), but have primarily focused on computational identification and have not verified their results using new antibodies that are not within the PDB training set. While there are many other structure prediction models other than AlphaFold2 (27, 28), including some specifically dedicated to predicting antibodies or antibody-like structures (29–32), we chose AlphaFold2 to directly test its ability to correctly identify and place epitopes into an antibody binding cleft. We selected AlphaFold2 due to its widespread use throughout the literature, as well as its ease of installation and modification via the LocalColabFold implementation (23). Another reason for selecting AF2 is to attempt to quantify its abilities the compare simple linear epitopes, since the team behind AF-multimer reported that conformational antibody complexes were difficult to predict accurately (12). In this project we test a method we call PAbFold, a LocalColabFold-based pipeline to identify epitopes for several well-known linear-epitope antibodies from sequence information only. There was a strong correlation between AlphaFold2’s confidence in the peptide structure (pLDDT) (33) and the experimentally verified epitope binding sequence. Additionally, we found that AlphaFold2 very accurately predicted the linear epitope of a novel SARS-CoV-2 nucleocapsid-specific antibody (mBG17) with minimal prior epitope information. The molecular interactions predicted by AlphaFold2 were experimentally validated using peptide mapping ELISA experiments. Overall, this work demonstrates that AlphaFold2 has compelling promise for linear antibody epitope discovery from sequence information alone. We also have observed that this emergent linear epitope prediction ability is sensitive to the peptide length and that the performance was optimal when using AlphaFold2-multimer version 2 and older MSAs generated by MMSEQS version 2202 server, rather than the more recent AlphaFold2-multimer version 3 models and MMSEQS version 2302 server.

## Materials and Methods

### Software

All structure predictions were completed on a single AMD EPYC 7443 server with two NVIDIA RTX A5000 GPU cards. PAbFold code was written in Python 3.7 and Bash. The only extra Python dependencies are NumPy and Matplotlib. AlphaFold2 calculations were run using an installation of LocalColabFold (23). Briefly, PAbFold contains 3 stages. In the first stage, a python script ‘A_PeptideMapping_prep_submission_files.py’ writes FASTA input files for ColabFold. Each FASTA file contains the entire sequence of the subject scFv, a colon “:”, and then the candidate linear epitope which represents a small section of the target antigen protein that changes dependent upon both the epitope length (default 10 a.a.) and a sliding window (default 1 a.a.).

After completion of the ColabFold jobs, two different analysis methods are presented in this paper, and both are accessible via the ‘B_PeptideMapping_plddt_perres_analysis.py’ python script. The first is the ‘Simple Max’ method, which assesses each peptide window with only the output model that is top ranked by ColabFold (on the basis of ipTM). The AlphaFold2 confidence pLDDT (33) is recorded for each residue within the peptide. Other than the N- and C-terminal residues, each residue is observed within multiple windows. We proceed to calculate (and plot) the maximum pLDDT observed for each residue across the set of sliding window peptides that contain that residue. Thus, in the ‘Simple Max’ method each residue is considered independently. To obtain aggregate scores for each peptide window, we sum the maximum pLDDT associated with each member residue. This method is sensitive in that any isolated high-confidence residue placements in the top ranked AlphaFold2 peptide prediction can increase the score, but a high aggregate peptide score could arise from multiple, mutually inconsistent peptide binding poses. Our second, complementary analysis method instead focuses on recognizing full peptide poses of elevated AlphaFold2 confidence. We refer to the second method as the ‘Consensus method’ because it begins by averaging the per-residue pLDDT across the five AlphaFold2 models. We then compute the average pLDDT for each peptide. For visual inspection, scripts output a heat map for the average per-residue pLDDT and a bar-chart that for the subsequent per-peptide average pLDDT. In this case, we simply rank top peptides based on the per-peptide average pLDDT. Scripts are available at https://github.com/jbderoo/PAbFold.

### Antibody sequences

Sequences and references for antibodies, scFvs, and antigens can be found in **Supplemental Table 1A**. To create an scFv, the complementarity determining regions or loops of an antibody are identified via the Kabat numbering scheme. The loops are then spliced onto the scFv backbones of the 15F11 and 2E2 as previously described by our group (22). The scFv sequences are aligned with their CDR loops and flexible linkers highlighted in **Supplemental Table 1B**.

### Monoclonal Antibody Production

Anti-SARS-CoV-2 nucleocapsid protein (NP) monoclonal mouse antibody mBG17 was previously developed and characterized (34). Briefly, two BALB/c mice immunized with recombinant NP were sacrificed and primary splenocytes isolated. Splenocytes were fused with Sp2/0 Ag14 myeloma cells and individual hybridoma clones were isolated after eleven days. Hybridoma clones were tested for antibody production against NP via enzyme-linked immunosorbent assay (ELISA) and western blot. Clones were further tested for isotype and cross-reactivity, and VH and VL sequences were determined. The hybridoma clone mBG17 was identified as a SARS-CoV-2 nucleocapsid-specific antibody targeting linear epitope via ELISA and western blot (34). Generation of recombinant mBG17 and production of recombinant antibody in 293F cells was previously described (34). The approximate epitope region for mBG17 was determined via western blot with modified recombinant NP proteins containing 40 to 50 amino acid deletions. The epitope location was determined to reside between SARS-CoV-2 nucleocapsid residues a.a. 381-419 based on loss of western blot signal with the a.a. 381-419 deletion (34).

### Peptide Competition ELISA

The anti-SARS-CoV-2 nucleocapsid protein mBG17 antibody epitope was experimentally identified using competition enzyme-linked immunosorbent assay (ELISA). Using the previously determined 39 nucleocapsid protein amino acid range for the mBG17 epitope as a starting point, seven overlapping peptides were synthesized (Thermo Scientific) spanning the 39 amino acid region with overlaps of 5 amino. These peptides were termed Fragment 1 through 7 (**Table 1**). A 96-well ELISA plate was coated with 0.1ug/ml of recombinant SARS-CoV-2 NP (34) overnight at 4°C. The plate was blocked with 4% (w/v) dry non-fat milk in 1X PBS with 0.1% (v/v) Tween-20 for 1 h shaking at room temperature. While blocking, inhibited recombinant mBG17 antibody samples were produced by incubating 40 μL of antibody with 40 μg (approximately 30 nMol) of a single peptide fragment for one hour at room temperature. Following this, peptide-incubated mBG17 was applied to the blocked nucleocapsid protein coated plate in triplicate and allowed to incubate for 1 h at room temperature while shaking. The plate was rinsed with 0.1% (v/v) Tween-20 in 1X PBS and washed three more times for 5 minutes shaking at room temperature. The plate was then incubated with HRP-conjugated goat anti-mouse polyclonal antibody solution diluted at 1:20,000 in 1X PBS for 1 h shaking at room temperature. After another rinse and three more washes the plate was developed with 1-Step™ Ultra TMB-ELISA Solution (ThermoFisher) before stopping the reaction with an equal volume of 2M H2SO4. Solution absorbance at 450 nm was measured using a PerkinElmer Victor X5 multilabel plate reader. Absorbances were averaged within fragment-inhibited sample groups and corrected with the average value of the negative control. These absorbances were then normalized against the absorbance from the group with the highest value before multiplying by 100 to obtain percentage of potential signal.

**Table 1:**
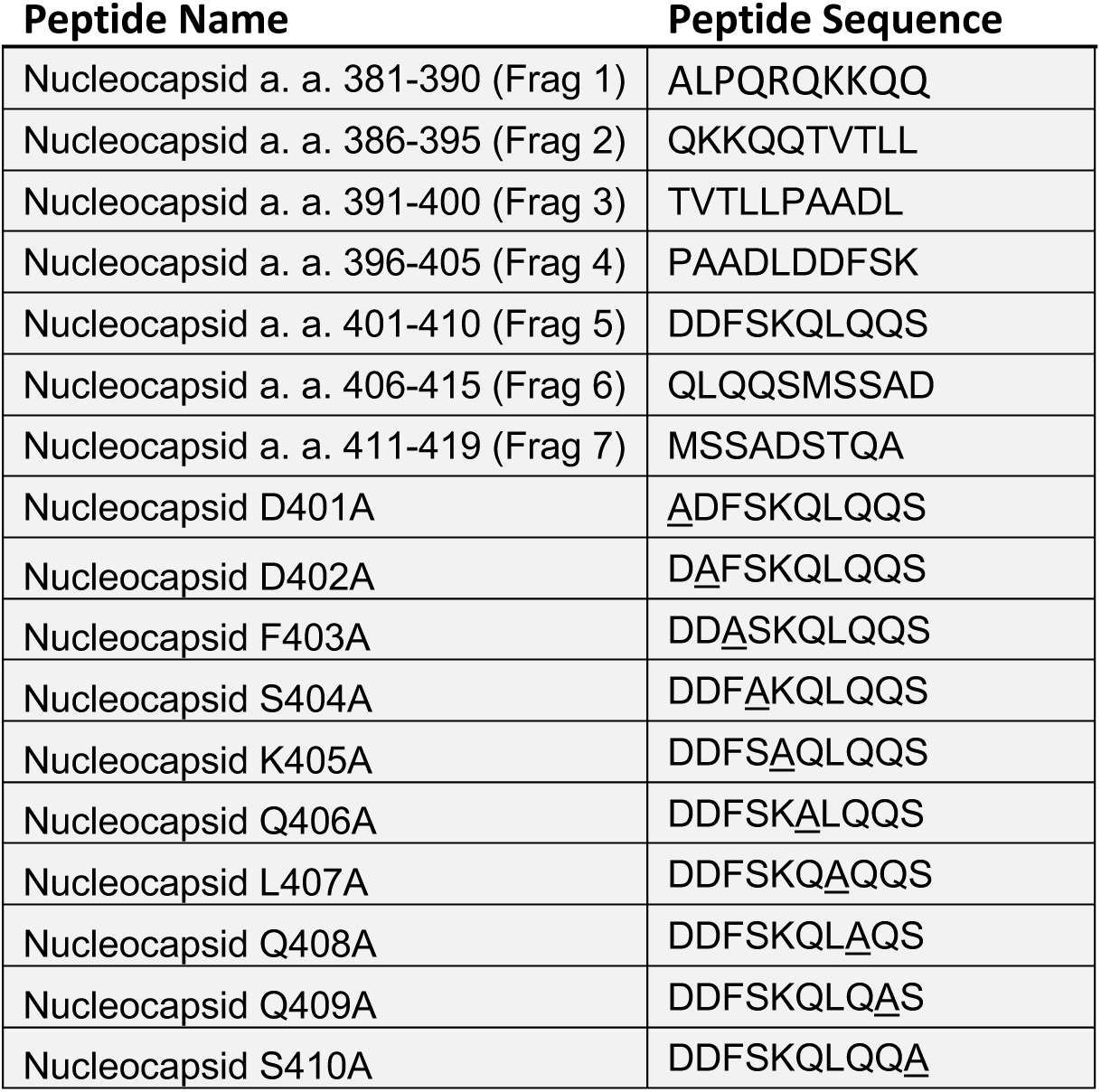

The effect of single alanine substitutions on fragment 5 (DDFSKQLQQS) peptide binding was determined by competition ELISA using a series of ten alanine-substituted peptides (**Table 1**) at a range of concentrations to determine relative competition activity. A modified version of the previously described inhibition ELISA was performed using the unmodified Fragment 5 peptide and the ten alanine-substituted peptides. During the mBG17 inhibition step, the mBG17 antibody solution was incubated with a 4-fold serial dilution of peptides beginning at 40 μg and continuing to ∼2.5 ng before being applied to the NP coated plates in triplicate. The remainder of the competition ELISA was carried out as described above.

### Assessment of AlphaFold2 generated scFv structures

We first verified that AlphaFold2 could generate scFv structures that have similar structures to their parent monoclonal antibodies. We chose the 9E10 clone of the anti-Myc antibody as an initial test system, as the scFv sequence is available (35) and has a well-known linear epitope (EQKLISEEDL)(36). We predicted the wild-type Myc scFv structure and aligned this model to the corresponding Fab crystal structure (PDB entry 2orb) via the align command in PyMOL (**Supplemental Figure 1A**). The AlphaFold2 predicted scFv was very similar (RMSD value of 0.42Å) to the anti-Myc Fab structure, suggesting that the predicted scFv structure was a suitable starting point for epitope prediction. We also examined the structures of the Myc CDRs loop grafted onto the 15F11 (37) and 2E2 (22) frameworks, as we have previously observed that loop grafting onto these frameworks can enhance protein folding and solubility (22). The loop-grafted Myc-2E2 and Myc-15F11 and structures were also similar to the Myc Fab structure (PDB 2ORB) (36) with similar RMSD values of 0.45Å (**Supplemental Figure 1B**), indicating that they are also reasonable starting points for epitope prediction.

## Results

### Development of Python-based scripts for automated scFv:peptide structure prediction

We developed a series of Python scripts that automate the process of epitope prediction and analysis with AF2. A_Peptide_Mapping_prep_submission_files.py accepts a linear scFv sequence and a linear full-length antigen sequence, and processes the antigen sequence into a series of short peptides with custom peptide length and sliding window sizes (default parameters are 10 amino acid peptides with a 1 amino acid sliding window). It then adds lines for each scFv:peptide pair to a FASTA file. Structures are then predicted via LocalColabFold for each scFv:peptide pair with AlphaFold2 in parallel on two NVIDIA RTX A5000 GPUs. The python script B_PeptideMapping_plddt_perres_analysis.py parses the AlphaFold2 output structures to extract per-residue pLDDT for the peptide residues in each scFv:peptide pair. Conf_plot_and_top10.py will plot the maximum pLDDT (across all host peptides) scores as a function of amino acid position within the antigen sequence and ranks predicted peptides based on ΣpLDDT scores for the ‘Simple max’ method. To use the ‘Consensus’ method, include the –all-models flag when running B_PeptideMapping_plddt_perres_analysis.py. We also supply a python script that replicates how we present the data called all_model_analysis.py for use.

An overview of the method is shown in **Figure 1**. AF2’s failure to predict whole antigen structure coupled with the scFv is highlighted in **Supplemental Figure 2**. Both the ‘Simple Max’ and ‘Consensus’ methods were calculated first by parsing every pLDDT score received by every residue in the antigen sequence sliding window output structures. From the resulting data structure, the Simple Max method simply finds the maximum pLDDT value ever seen for a single residue (across all sliding windows and AF2 models). For the Consensus method, per-residue pLDDT was first averaged across the 5 AF2 models. These averages are reported in the heatmap view and further averaged per sliding window for the bar chart below. In principle, the strategy behind the Consensus method is to take into account agreement across the 5 AF2 models and provide insight into the confidence of entire epitopes (whole sliding windows of n=10 default) instead of disconnected, per-residue pLDDT maxima. Having two scoring metrics is useful because the selection of predicted hits can differ. As shown in Figure 2, part of the Myc epitope makes it into the top 5 peptides when selection is based on summing per-residue maximum pLDDT (despite there being no requirement that these values originate in the same physical prediction). In contrast, a Consensus method score more directly reports on a specific sliding window, and the strength of the highest confidence peptides is more directly revealed with superior signal to noise as shown in Figure 3. Variability in the ranking of top hits between the two methods arises from the fundamental difference in strategy (peptide-centric or residue-centric scoring) as well as close competition between the raw AF2 confidence in the known peptide and competing decoy sequences.

**Figure 1.**
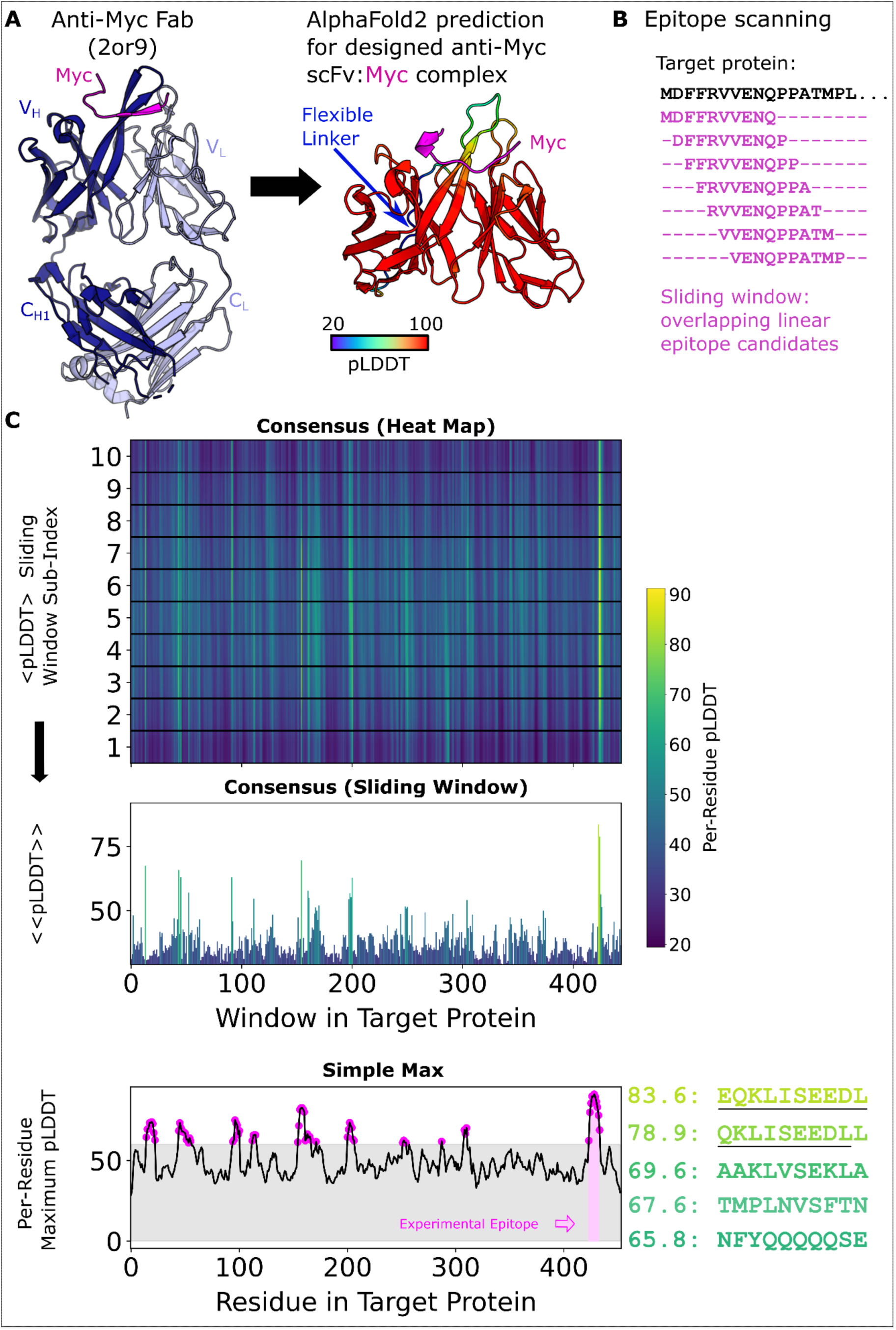
PAbFold pipeline for linear epitope prediction. **A)** Antibody V_H_ and V_L_ protein sequences are used to generate scFv sequences, either based on the native antibody sequences or loop grafting complementarity determining regions (CDRs) onto either the 2E2 or 15F11 antibody framework regions (2E2 shown). **B)** The target antigen sequence is parsed into a list of small overlapping peptide sequences, with peptide step and window size parameters adjusted as needed. Rank ordered peptides are output, and partial epitope sequences are underlined manually to highlight the identification of the correct sequence. **C)** The scFv sequences from Panel A are co-folded with each of the peptide sequences derived from the target antigen in parallel batch mode on a GPU server. pLDDT scores from each structure prediction experiment are collected and scores are presented in their sliding window, both as a heat map organized along the length of the target antigen sequence and a bar chart that shows the per-peptide average pLDDT (Consensus Method). Additionally, the Simple Max data is presented in the third and final panel.

**Figure 2.**
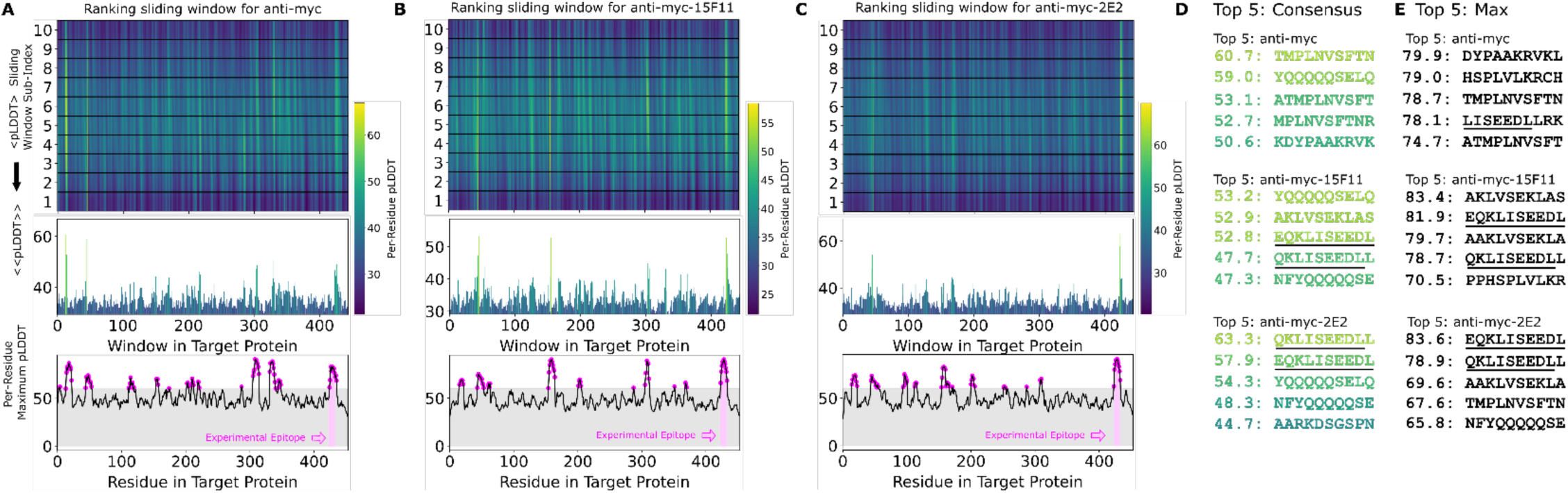
The Alphafold2-based PAbFold method predicted the Myc linear epitope in different scFv backbones. The anti-Myc V_H_ and V_L_ antibody sequences were used to generate either **A)** wild-type Myc scFv or loop grafted chimeric **B)** Myc-15F11 or **C)** Myc-2E2 scFv variants. The Myc proto-oncogene protein sequence (Genbank NP_001341799.1) was used as the target antigen and processed into 10 amino acid overlapping peptides with a 1 amino acid sliding window. The structure for each scFv:peptide pair was predicted with AlphaFold2 in batch mode on two NVIDIA A5000 GPUs. Average consensus pLDDT values for each scFv:peptide window are illustrated, as well as the maximum pLDDT observed for each residue in any window (bottom). **D)** Top ranking binding peptides based on average consensus pLDDT. **E)** Top ranked binding peptides based on summing per-residue maximum pLDDT. For D and E, underlining represents overlap with the reported Myc epitope (EQKLISEEDL).

**Figure 3:**
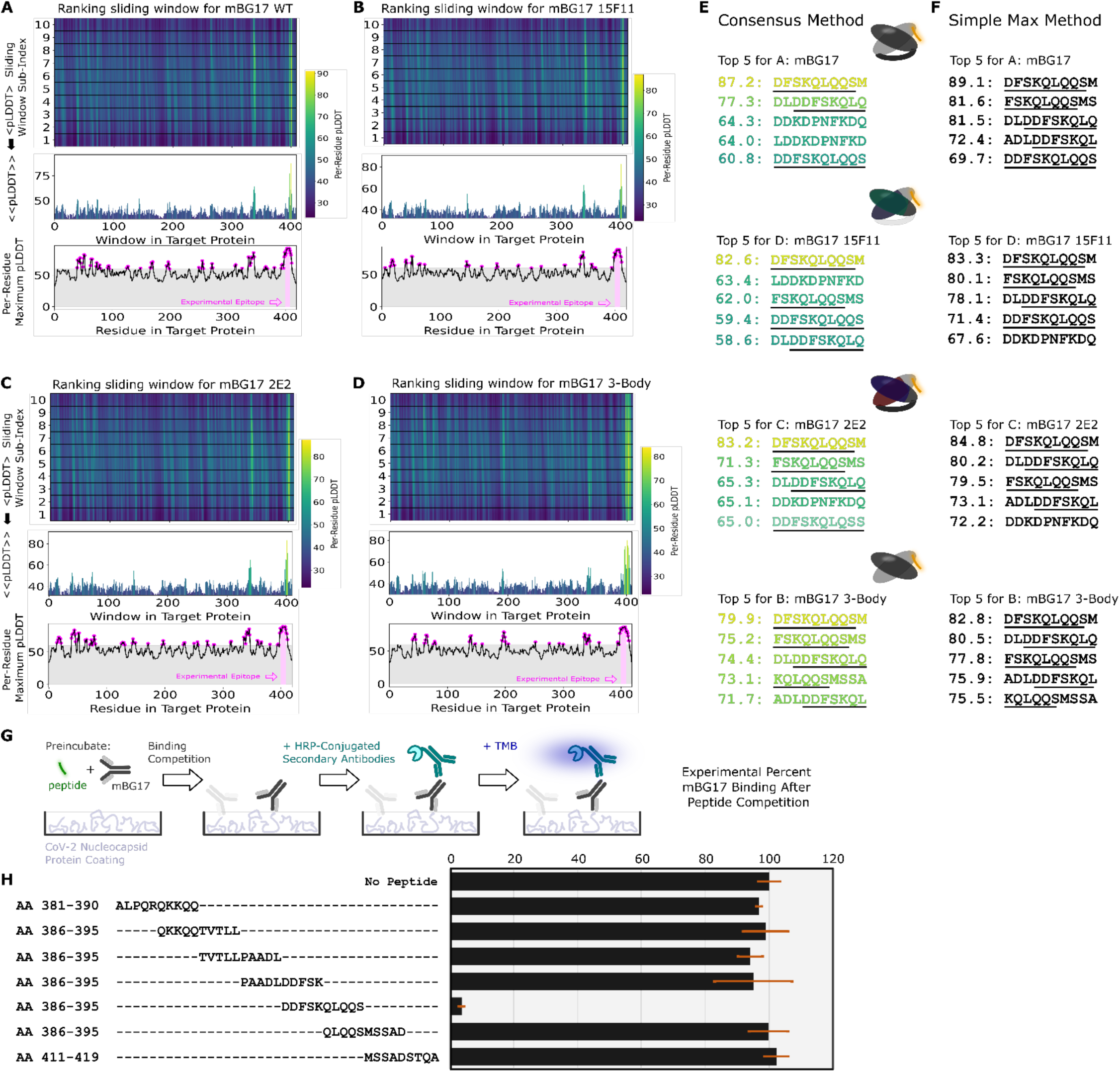
The AlphaFold2-driven PAbFold epitope scan method can accurately identify a linear epitope for a novel SARS-CoV-2 antibody. Antibody VH and VL sequences for SARS-CoV-2 nucleocapsid protein targeted antibody were used to generate scFv sequences **A)** WT, **B)** 15F11, **C)** 2E2 or native VH and VL sequences **D)** 3 body). Variant scFv sequence in complex with peptide windows from the SARS-CoV-2 nucleocapsid protein (Genbank Accession: YP_009724397) were subjected to AlphaFold2 structure prediction. The top 5 peptides ranked by either the **E)** Consensus method or the **F)** Simple Max method, with the underlined sequence highlighting the experimentally verified sequences and a cartoon schematic for each system shown. **G)** Competition ELISA schematic for assessing the ability of synthetic peptides derived from the SARS-CoV-2 nucleocapsid protein. **H)** Amino acid windows showing binding interference, with mBG17 binding to SARS-CoV-2 nucleocapsid protein (n = 3). Percentage of binding values were calculated from the no-peptide control. Alignment of synthetic peptides corresponding to SARS-CoV-2 nucleocapsid a. a. 381-419. Peptide a. a. 401-410, which demonstrated mBG17 competition.

### Testing of scFv:peptide structure prediction method using the Myc Epitope

We first tested the PAbFold method with the anti-Myc-scFv described in (38), using the full-length human Myc proto-oncogene protein sequence as the antigen. We initially used an antigen peptide length of 10 and a 1 amino acid sliding window. Given these parameters, the 9 a.a. Myc epitope motif (EQKLISEEDL) appeared intact within one of the 10-mer peptides, with subsets of the 8, 9, 11, and 12 a.a. appearing in neighboring sliding peptide windows. PAbFold generated predicted structures, each of which took an average of ∼200 seconds to process. The entire process took approximately 12 hours on our GPU server. AlphaFold2 placed all peptides into or near the traditional antigen binding site between the CDR loops (**Supplemental Figure 3**). The average confidence (mean pLDDT across residues) for these peptides ranged from 20 to 90. When we inspect the consensus confidence for each residue in each sliding window (**Figure 2A**), the expected Myc peptide epitope (EQKLISEEDL) was one of several peptides with high average pLDDT. The second highest ranked peptide in this analysis (QKLISEEDLL) was a near perfect match for the expected epitope. We consider this window to be a successful prediction. Perhaps surprisingly, the peptide window with the exact match (EQKLISEEDL) did not score particularly well due to its average pLDDT of 51.0. In this instance, the expected epitope sequence did not stand out when plotting the maximum observed per-residue pLDDT for each residue (**Figure 2A, bottom**).

We proceeded to test predictions with two engineered scFv chimeras where loop grafting was used to place the Myc recognition CDRs onto two antibody framework regions with high *in vivo* performance, generating Myc-15F11 and Myc-2E2 scFv sequences. Epitope prediction performance was markedly improved with the chimeric scFvs (**Figure 2B and 2C**). Specifically, the QKLISEEDLL peptide window became the top ranked peptide on the basis of average consensus pLDDT. In the case of Myc-2E2 (**Figure 2C**), the average confidence for the correctly predicted epitope was particularly high compared to alternate peptide windows, and another close match to the expected epitope (EEQKLISEED) was ranked within the top 5 peptides (**Figure 2D**). Ranking epitopes using the Simple Max analysis was similar; the region containing the correct epitope was nearly top ranked for Myc-15F11 and was top ranked for Myc-2E2 (**Figure 2E**). Thus, AlphaFold2 was able to more clearly detect authentic Myc antibody epitope using CDRs loop grafted onto the 2E2 or 15F11 frameworks, relative to the native Myc scFv framework.

To investigate the superior epitope recognition performance of the chimeric Myc scFvs, we aligned the Cα coordinates for the predicted scFv structures (predicted with and without the target epitope) to the reference crystal structure and calculated the RMSD for all backbone positions (N, Cα, C, O) and the loops (**Supplemental Figure 4**). Notably, regardless of the Myc scFv variant, the CDR loop RMSD improved by more than 1Å when the epitope was present. Secondly, consistent with the improved epitope prediction performance for the chimeric scFvs (15F11 and 2E2), the epitope peptide QKLISEEDL was placed more accurately for those predicted structures than in the WT scFv (**Supplemental Figure 4**). We could not discern an obvious structural difference between the WT and chimeric scFvs that explains the structure prediction performance gap.

### Assessment of peptide length, sliding window size, and position on AlphaFold2 scFv:peptide structure prediction

Our initial selection of the 10 a.a. window was intended to match or slightly exceed the size of known epitopes such as Myc and HA. We next assessed how different peptide sizes and sliding window lengths would affect epitope prediction accuracy and run time. We re-ran the Myc-2E2-scFv:peptide complex prediction calculations varying peptide size between 8, 9, 10, and 11 (with a fixed sliding window size of 2) or varying the sliding window size to 1 or 5 (with a fixed peptide size of 10). We observed that using a sliding window of 2 a.a. provided nearly the same level of accuracy and resolution as the 1 a.a. Ultimately, we determined that our original peptide size of 10 amino acids and sliding window of 1 a.a. provided highest resolution data possible (**Supplemental Figure 5**) and therefore maintained a peptide size of 10 and a sliding window length of 1 for our remaining experiments.

We then predicted the complex structure for Myc-2E2 with various negative control peptides: A10, (GS)5, (GGGGS)2, and G10 to determine how non-binding peptides are docked and scored (**Supplemental Figure 5I and 5J**). We again observed that AlphaFold2 placed all peptides into the traditional antigen binding between the CDR loops, but the reported peptide scores for the negative controls were particularly low (29–41). These results indicate that AlphaFold2 “knows” where antigens bind in antibody or scFv structures and attempts to model any peptide partner into this region, but the low pLDDT scores indicate confidence in the interactions are quite low.

We also tested if AlphaFold2 could detect the Myc epitope if it was inserted as an epitope tag within different positions of a heterologous protein. We created a synthetic antigen by adding the Myc epitope within the 99-a.a. unrelated HIV-1 Gag protease protein sequence at either the N- or C-terminus or in the middle of the protein sequence, and used PAbFold to detect the Myc peptide (**Supplemental Figure 6**). In each case, the average consensus pLDDT was highest for the inserted epitope, such that the authentic epitope would be top ranked and prioritized for testing. Thus, as expected for a sliding window analysis, the epitope position within the antigen was no barrier to detection.

### Testing of the PAbFold method using the HA Epitope

Based on our success detecting the Myc epitope, we sought to determine if our method could detect a different well-known linear peptide, HA, derived from positions 114-126 within the Influenza A virus hemagglutinin protein (YDVPDYASLR). Using an anti-HA scFv sequence that had been previously generated (22, 38), we generated new HA-15F11 and HA-2E2 scFvs loop grafted sequences. We used the same procedure described above to predict structures for influenza A virus HA derived peptides on HA-scFv (**Supplemental Figure 7A**), HA-15F11-scFv (**Supplemental Figure 7B**) and HA-2E2-scFv (**Supplemental Figure 7C**). In the HA case, the expected epitope was ranked highly for all three scFv variants, but when assessing entire peptides by average consensus pLDDT was only ranked in the top 5 for the HA-15F11-scFv. These results, in combination with the Myc results described above, indicate that AlphaFold2 can accurately detect linear antibody epitopes in antigen sequences, and that grafting CDR loops onto alternative scFv backbones may increase the noise-to-signal ratio, making the identification of correct epitopes more accurate.

Like the Myc system, trends are observed with the HA system regarding loop placement. Although not as extreme, the loops for all HA scFvs undergo movement that make it more closely match the crystal structure (PDB entry 1frg). Again, the epitope placement of predicted structures of the chimeric scFvs more closely mimicked the deposited crystal structure than the WT scFv (**Supplemental Figure 4B**).

### Determination and experimental validation of a novel linear antibody epitope

The Myc and HA monoclonal antibodies are well known and several crystal structures (Myc PDB: 2or9, peptide bound (2009) | HA PDB:1frg, peptide bound (1994)) have been solved (22, 36, 38, 39), raising the possibility that AlphaFold2 has incorporated these antibody or epitope structures into its training set. The AlphaFold2 training set was reported to exclude chains of less than 10, which would eliminate the myc and HA epitope peptides. Nonetheless, to guard against the possibility that the AlphaFold2 models have incorporated specific knowledge into the training set thereby directly probing if PAbFold epitope scanning can predict a linear antibody epitope without *a priori* knowledge of the antibody or antigen sequence, we tested if PAbFold can predict the epitope sequence of a recently developed antibody lacking structural information available in the Protein Data Bank. The mBG17 mouse monoclonal antibody was generated in response to the COVID-19 pandemic, the antibody VH and VL sequences were determined, and the epitope was localized to a. a. 381-419 via Western blot analysis of deletion mutants of the nucleocapsid protein (34). mBG17 was not included in AlphaFold2’s training or test set, making it an ideal test case for *de novo* epitope prediction.

The mBG17 monoclonal antibody was converted to wild-type scFv, 15F11-scFv, and 2E2-scFv using the same procedures used for Myc and HA scFv. As an additional control calculation (labeled “3-body”), we used AlphaFold2 to predict the structure for a 3-protein complex (the peptide, and the disconnected nontruncated mBG17 VH and VL variable domain sequences). All 4 Fab variants (WT scFv mBG17, 15F11-mBG17 scFv, 2E2-mBG17 scFv, and 3-body mBG17) were screened against all 10 a.a. peptides with a 1 a.a. sliding window, as with Myc and HA. In all 4 cases, AlphaFold2 predicted that the top ranked peptides were located in the a.a. 381-419 region of the SARS-CoV-2 nucleocapsid protein, and more specifically residues a.a. 400-415 (**Figure 3A**, **3B**, **3C, and 3D**). The top scoring peptide for all three scFv variants was the 402-411 window (DFSKQLQQSM) (**Figure 3E and 3F**). The strong AF2 preference for peptides from this C-terminal segment was particularly evident in the average consensus pLDDT analysis.

We next sought to experimentally verify the minimal linear epitope for mBG17 to determine how closely the AlphaFold2 prediction corresponded to our experimental data. Seven 10 a.a. peptides that overlapped by 5 a.a. each were synthesized and used in competition ELISAs with mBG17 monoclonal antibody and recombinant SARS-CoV-2 nucleocapsid protein (**Figure 3G and 3H**). The peptide corresponding to a.a. 401-410 showed almost complete competition of mBG17 binding to the SARS-CoV-2 nucleocapsid protein in the ELISA, whereas none of the other peptides were able to compete for mBG17 binding to nucleocapsid. Peptides a.a. 296-405 and a.a. 406-415 overlap a.a. 401-410 at the N- and C-terminus, respectively, but neither was able to compete, indicating that mBG17 binds a.a. 401-410 on both sides of a.a. 405 and a.a. 406. An alignment of all the peptides used in the overlapping peptide competition ELISA experiments showed that peptide sequence DDFSKQLQQS represents the experimentally determined epitope for mBG17, nearly identical to the epitope predicted by AlphaFold2 (**Figure 3H**: DDFSKQLQQS). These results demonstrate that the PAbFold pipeline was able to very accurately predict the region that an antibody binds to a novel linear epitope that is not present in AlphaFold2’s training set.

### Fine-characterization of the mBG17 epitope and comparison to the predicted AlphaFold2 model

To further experimentally characterize the binding of the mBG17 to the a.a. 401-410 (DDFSKQLQQS) peptide and compare experimental data with the predicted AlphaFold2 model, we designed and synthesized ten additional peptides, each containing an alanine point mutation at one position in the a.a. 401-410 peptide. The peptides are labeled D1A, D2A, F3A, S4A, K5A, Q6A, L7A, Q8A, Q9A, and S10A. Competition ELISAs were performed using increasing concentrations of each peptide to better assess differential binding (**Figure 4A**). As expected, WT (a.a. 401-410) peptide showed strong competition, although Q9A showed slightly better competition. This could be attributed to alanine’s propensity to be in an alpha-helical coil (PropA, AHC = 0) vs glutamine’s propensity to escape it (PropQ, AHC = 0.39) (40), thus further stabilizing the Q9A alpha helix. D1A showed no change in competition, indicating that D1 was not involved in binding. Peptides with substitutions K5A, Q6A, and S10A showed minor reductions in competition, S4A showed a moderate reduction on competition, whereas resides D2A, F3A, L7A, and Q8A all showed strong reductions in competition. These data indicate that the key interactions between mBG17 and the a.a. 401-410 peptide are residues D2, F3, L7, and Q8, with S4 playing a moderate role and D1, K5, Q6, Q9 and S10 playing negligible roles in binding.

**Figure 4.**
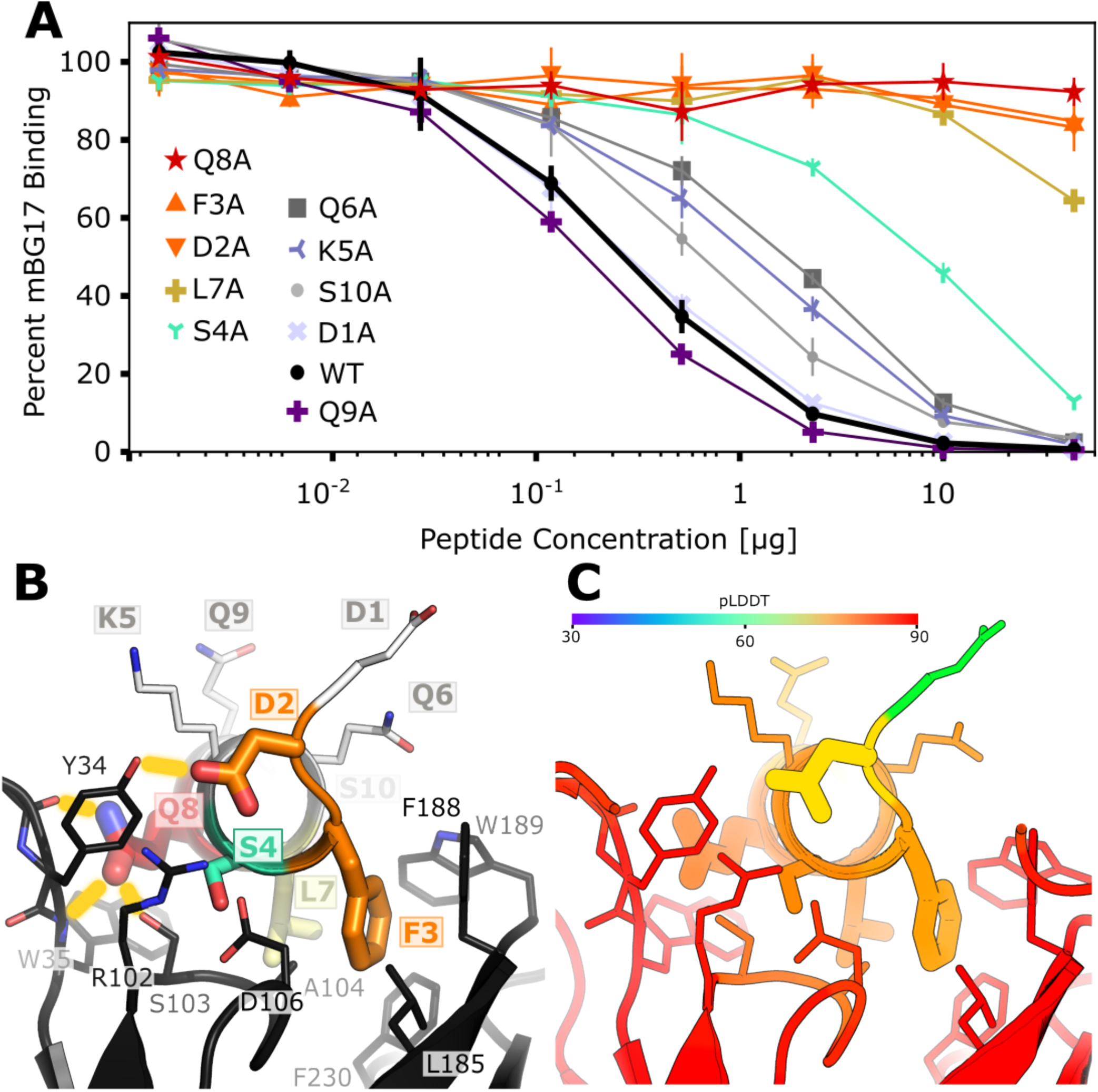
The Alphafold2-Driven PAbFold method accurately predicts molecular interactions between a linear epitope and a scFv. **A)** Competition ELISA assessing the ability of synthetic alanine mutant peptides derived from the SARS-CoV-2 nucleocapsid protein (a. a. 401-410: DDFSKQLQQS) to interfere with mBG17 binding to SARS-CoV-2 nucleocapsid protein (n = 3). Percentage of binding values were calculated from the no-peptide control. **B)** AlphaFold2 model for mBG17-15F11 scFv bound to a. a. 401-410 peptide (the average peptide pLDDT was 83.5). Residues that display sharply reduced binding to mBG17 upon mutation to alanine in competition ELISAs (D2, F3, S4, L7, Q8) are shown as warm-colored thick sticks. Predicted hydrogen bonds between the peptide and the scFv are depicted by yellow bars. Sites where mutation to alanine was less disruptive to binding (Q6A, K5A, S10A, D1A, and Q9A) are depicted as thin sticks with cool colors. The carbon atoms of residues in panel B are colored according to the corresponding data in panel A. **C)** The same AlphaFold2 model for the mBG17-15F11 scFv bound to a.a. 401-410 colored with confidence (pLDDT) as predicted by AF2.

Finally, we compared the experimental data shown above with the best scoring mBG17:DDFSKQLQQ model generated by AlphaFold2 (**Figure 4B and 4C**). The AlphaFold2 model suggests that residue D2 forms a hydrogen bond with mBG17 a.a. Y34, residue F3 forms a hydrophobic interaction with mBG17 a.a. L185, residue S4 lacks a hydrogen bond partner, residue L7 forms a hydrophobic interaction at the base of the binding cleft with mBG17 a.a. A104, and residue Q8 hydrogen bonds with the backbone carbonyl of Y34 and the backbone amide of W35. Residues that experimentally showed no or minimal effects on competition (D1, K5, Q6, Q9) are all predicted to interact primarily with the solvent and lacked visible interactions between the peptide and scFv sequence. In summary, the AlphaFold2-driven PAbFold prediction was remarkably consistent with the experimental alanine scanning data, suggesting that the prediction of the mBG17 linear epitope location was accurate due to the correct prediction of the structural details for how that linear epitope binds to the antibody.

## Discussion

In this project we assessed the ability of an AlphaFold2-based linear epitope scan pipeline we call PAbFold (Peptide:Antibody Fold) to predict linear antibody epitopes using just antibody and antigen sequences. We first assessed the quality of scFv models produced by AlphaFold2. We then developed a series of Python scripts that accept scFv and whole antigen protein sequences as inputs, parse the antigen protein sequences into short overlapping peptides, run batch predictions for each scFv:peptide pair, and output two peptide scoring schemes based on the peptide per-residue pLDDT scores as a metric for AlphaFold2 model confidence.

Binding of the expected epitope to the WT-Myc scFv could only be detected via the consensus method, but either analysis method could readily detect the expected epitope bound to the chimeric Myc scFvs. Conversely, the alternate analysis method (Simple Max) performed better with respect to ranking the expected HA epitope binding to the WT and chimeric anti-HA scFv variants. In the HA case, performance was comparable for both the WT and chimeric scFv variants.

It is important to note that binding of scFv variants to sequences other than the expected epitopes may be statistically unlikely but not impossible. For example, consider the peptide ATMPLNVSFT near the N-terminus of the Myc proto-oncogene protein sequence. In the context of the WT anti-Myc scFv this peptide had slightly higher average consensus pLDDT (52.4 rather than 51.0) than a peptide (QKLISEEDLL) that closely matched the expected epitope. In the absence of direct experimental evidence, predicted affinity for this unexpected sequence is not necessarily incorrect, though the lack of comparable predicted binding to the 15F11 and 2E2 chimeric scFv variants further decreases the likelihood. In the future, it might be useful to assess peptide binding via consensus across scFv variants.

Lastly, we tested this process on a novel antibody generated by our group targeting the SARS-CoV-2 nucleocapsid protein (mBG17) and found the method performed significantly better than with Myc and HA. Either analysis method could very easily flag peptide windows containing the authentic experimentally validated epitope. This worked for the WT scFv, the chimeric scFv variants, and even a structure with disconnected heavy and light chain domains. Experimentally, we cleanly validated the AlphaFold2 prediction using a peptide competition ELISA assay to experimentally determine the mBG17 epitope. Confidence in the AlphaFold2 prediction was further buoyed via alanine scanning peptide competition ELISAs that verified the importance of the key binding interactions predicted by AlphaFold2.

Identification of antibody VH and VL sequences from monoclonal B-cells has become a routine task, with sequence information obtainable via various sequencing technologies such as next generation sequencing and nanopore sequencing for a relatively low cost. As a result, the determination of the epitope in service of a deeper understanding of how antibodies bind their antigen is an increasingly notable bottleneck. An experimental epitope determination campaign can take weeks or months of work, but with the advent of AlphaFold2 and the epitope prediction method we describe here, an antibody and its antigen could be sequenced in a few days (often through contract research organizations for low cost) and accurate linear epitope predictions generated within less than a day, dramatically epitope validation throughput as well as providing detailed predictions for the molecular features of antibody-epitope interaction.

Conformational epitopes are structured antigens that are found during many immune responses, and prediction of these epitopes from antibody and antigen sequences would be a significant boon to the field of biology. For example, conformational epitope prediction coupled with single-cell B-cell sequencing would allow for detailed analysis of antibody maturation during immune responses to vaccines or pathogen infection, helping better define how the immune response to infection evolves over time and how evolution of antigen sequences affects the antibody response. In this work we did not focus on using AlphaFold2 to predict conformational epitopes primarily because of the complex structures that conformational epitopes possess. Literature reports suggest that prediction of the complexes between antibodies and both whole antigens and conformational epitope proteins has proven to be very difficult for AlphaFold2, and indeed the authors themselves make this observation (12, 41, 42). Notably, the structures that proved most difficult to predict for AF2 and other tools in the CASP15-CAPRI154 challenges were antibody-antigen complexes (43). Reports suggest that a mix of both statistics-based approaches (neural networks like AF2) and physics-based approaches (such as Rosetta) predict optimal antibody-antigen complexes (44). Indeed, if we attempt to predict binding of our scFvs to intact antigen proteins (**Supplemental Figure 2**), we find no predictive capability. When predicting scFv:peptide complexes, it may be the case that AlphaFold2 is able to thoroughly evaluate an induced fit for the peptide due to both its length (small sample space) and its propensity to not adopt a strong competing structure. In contrast, embedding the epitope within a larger and more complicated structure appears to degrade the ability of AlphaFold2 to sample a comparable bound structure within the allotted recycle steps. Additional complexities may arise in extreme induced conformational changes during docking. Recent reports indicate that progress is being made in predicting the binding locations of conformational epitopes (45, 46).

We observed that the ability of AlphaFold2 to successfully predict the epitope peptide binding is quite delicate. First, epitope prediction was highly sensitive to the peptide length (**Supplemental Figure 5**), with minimal predictive power for peptide length other than 10 a.a. Further investigation of this sensitivity would be a useful avenue for future research. Perhaps with enhanced sampling, epitopes can be detected within longer peptides (e.g. 11 a.a., 12 a.a., etc.). Methodological tuning of this type could ultimately help illuminate the path to increasingly difficult protein-protein binding prediction problems. Similarly, we have likewise determined that epitope scanning performance was sensitive to changes in the underlying AlphaFold2 neural networks and the MSA. Specifically, unless otherwise noted, all data in this report was obtained using ColabFold version 1.5.2 and the 5 neural networks that comprise AlphaFold2 multimer version 2 (mm2). Likewise, the MSAs we use were obtained from the MMSEQS server (and cached) when the default sequence databases were UniRef30 2202 and PDB70 220313. They have since been updated to PDB30 2302 and PDB100 230517. For a complete description, see the change logs on the github for ColabFold (https://github.com/sokrypton/ColabFold#colabfold---v152).

Insofar as protein-peptide prediction is an emergent “off-label” capability for AlphaFold2 that is not part of the training sets, further training of the models or other changes can degrade performance. Benchmarking performance can be difficult when there are multiple moving targets. The most recent calculations we have analyzed were using ColabFold version 1.5.2 which was current as of February 19, 2023. The changes from ColabFold 1.5.2 to 1.5.5 (current as of this writing) are limited to version control and ensuring ColabFold still works on Google Colab and therefore will not change the calculation performance. Relative to ColabFold 1.3 (the current method at the outset of this project), ColabFold 1.5.2 embodied two substantial changes. First, ColabFold 1.5.2 used the updated AlphaFold multimer (mm) version 3 by default. Second, the backend server MMSEQS ((47) and (https://github.com/soedinglab/MMseqs2)) that supplies MSAs also underwent updates, namely the database updates. Upon evaluation, we found that the recent default methods (ColabFold 1.5.2) still predicted the epitope successfully for the mBG17 system (**Supplemental Figure 8**). However, the ColabFold 1.5.2 default methods had a pronounced decline in PAbFold performance for the HA and Myc systems. Specifically, the combination of mm3 and the revamped ColabFold MSA server tended to be less discriminating compared to the default settings for ColabFold 1.3 (ColabFold 1.3 was the most up to date version when this project was initialized). The updated configuration flagged diverse peptide sequences with elevated pLDDT values (**Supplemental Figures 9 and 10**) resulting in the loss of successful epitope predictive power. While testing ColabFold 1.5.2 with the most recent MSA server, but reverting the AlphaFold2 models to mm2, the outcome improved, with experimentally validated sequences rising to the top more frequently than when using mm3 but still falling short in ranking the experimentally validated epitope sequence embedded within the antigen. However, when previously cached MSAs were paired with mm2 (using ColabFold 1.5.2), performance was maximized. Furthermore, we attempted to recreate the MSA databases locally with similar but not identical results to queueing the server with databases UniRef30 2202 and PDB70 220313 (**Supplemental Figure 11**). Additionally, the MMSEQS team ((47) and (https://github.com/soedinglab/MMseqs2)) graciously rebuilt a server we could query using LocalColabFold that mimicked the original UniRef30 2202 and PDB70 220313 database set up as closely as possible on their end. The MSA that was generated from these databases was used, and still did not perform as well as the original MSAs that were generated upon first retrieval and generation (**Supplemental Figure 12**). As a negative control, we repeated all calculations without using any MSAs and only relying upon the sequence to make a structural prediction. As expected, all epitopes were scored very poorly (**Supplemental Figure 13**). Despite our significant efforts, it is unclear why our initial results cannot be perfectly recapitulated, but the difference has been traced to detailed MSA contents (**Supplemental Figure 14**), resulting in differences in correct epitope identification. These results are summarized in (**Supplemental Figure 15**). These challenges are presumably compounded by the incredible diversity of the CDR loops in antibodies which could decrease the useful signal from the MSA as well as drive inconsistent MSA-dependent performance

One key lesson of this research effort is that caching the MSAs proved to be very useful as a method to guard against changes in the performance of 3^rd^ party tools. We recommend that future methods development work using LocalColabFold adopt the strategy of caching MSAs when feasible. It is also our hope that by describing the latent ability of AlphaFold2 to predict scFv-binding epitopes that this ability will be preserved and enhanced in future iterations.

## Acknowledgements

The authors would like to thank members of the Snow, Stasevich, and Geiss groups for helpful discussions at TagTeam meetings. This project was funded in part by NIH R01AI132668 (Geiss) and NIH R56AI155897 and R01AI168459 (Snow, Stasevich, Geiss). The authors would also like to thank Dr. Milot Mirdita for being our point of contact for the ColabFold and MMSEQS teams and assisting us with setting up databases and generating MSAs for our use.

## Supporting Information

**Supplemental Table 1A.**
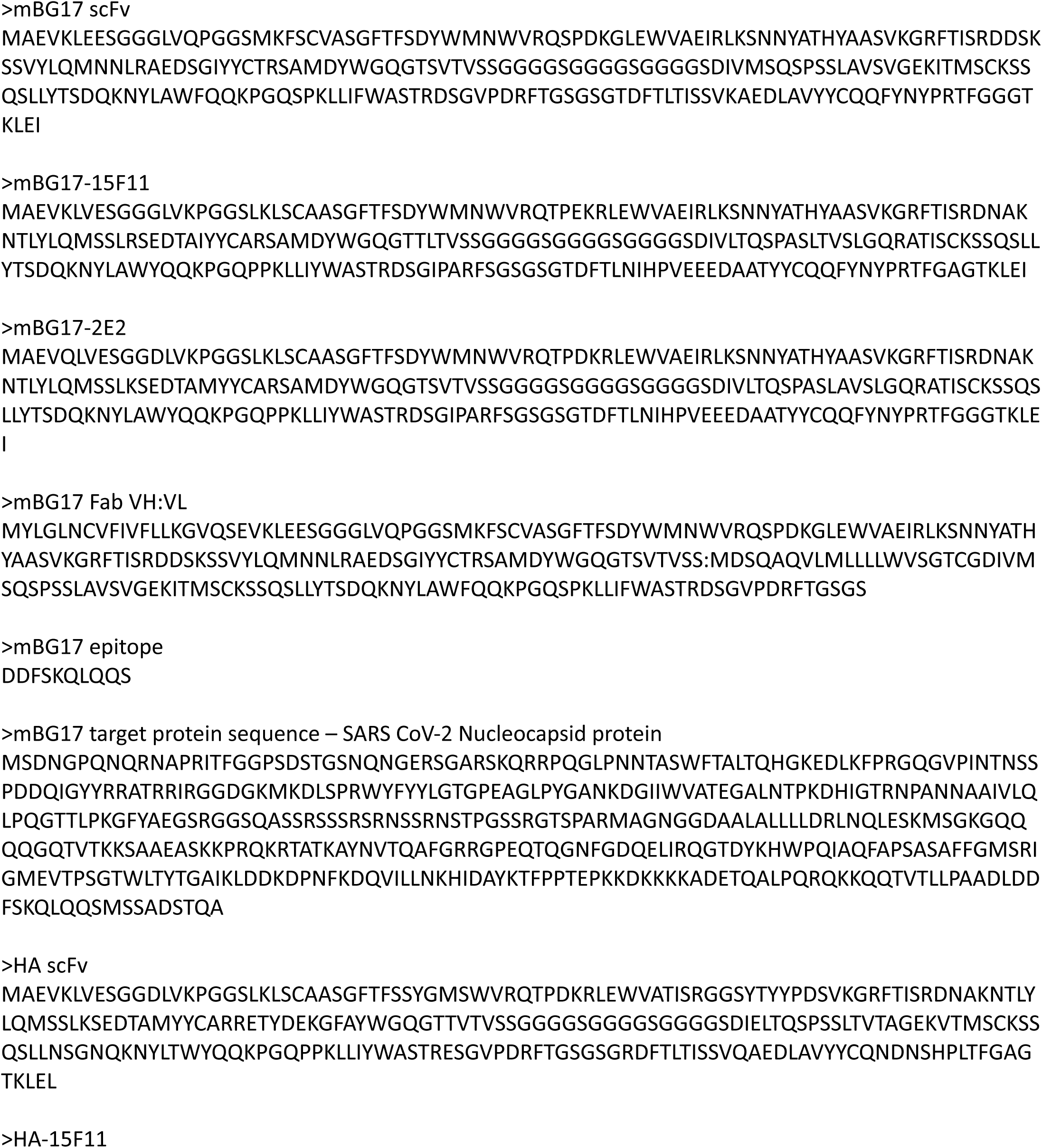
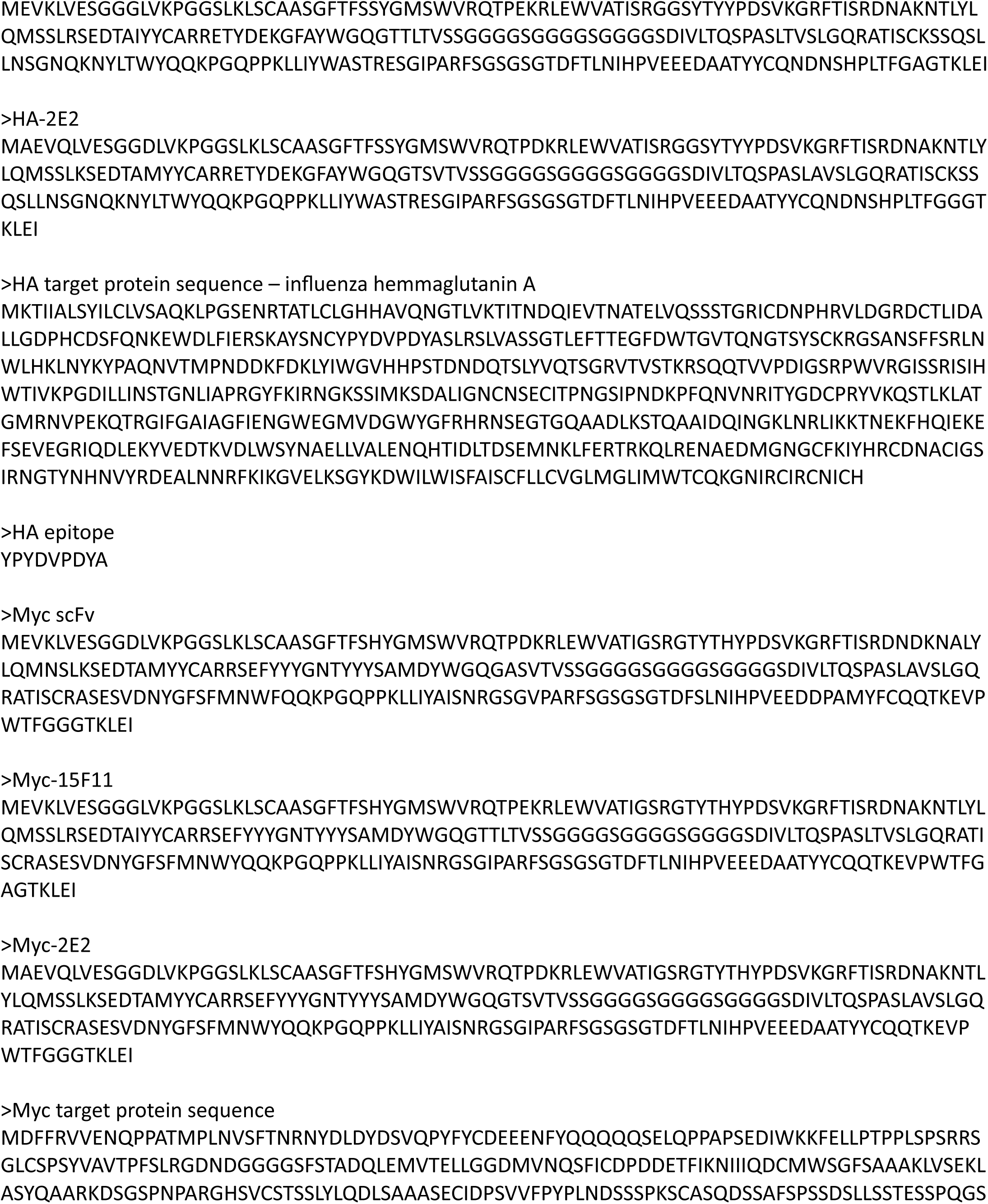
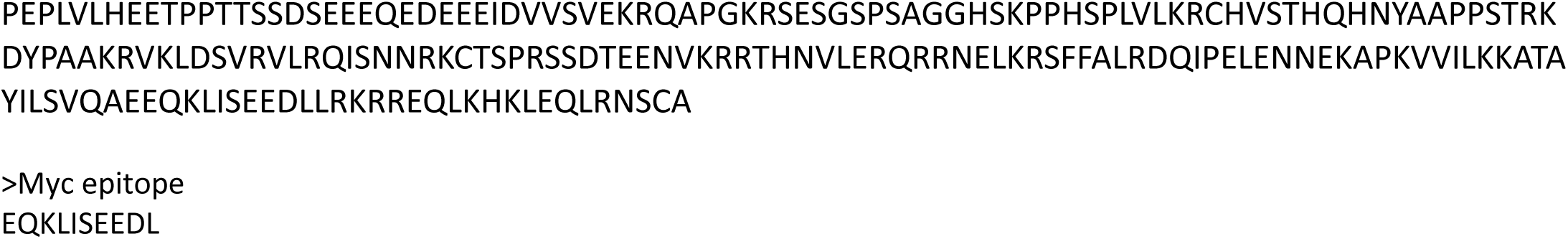

**Supplemental Table 1B.**
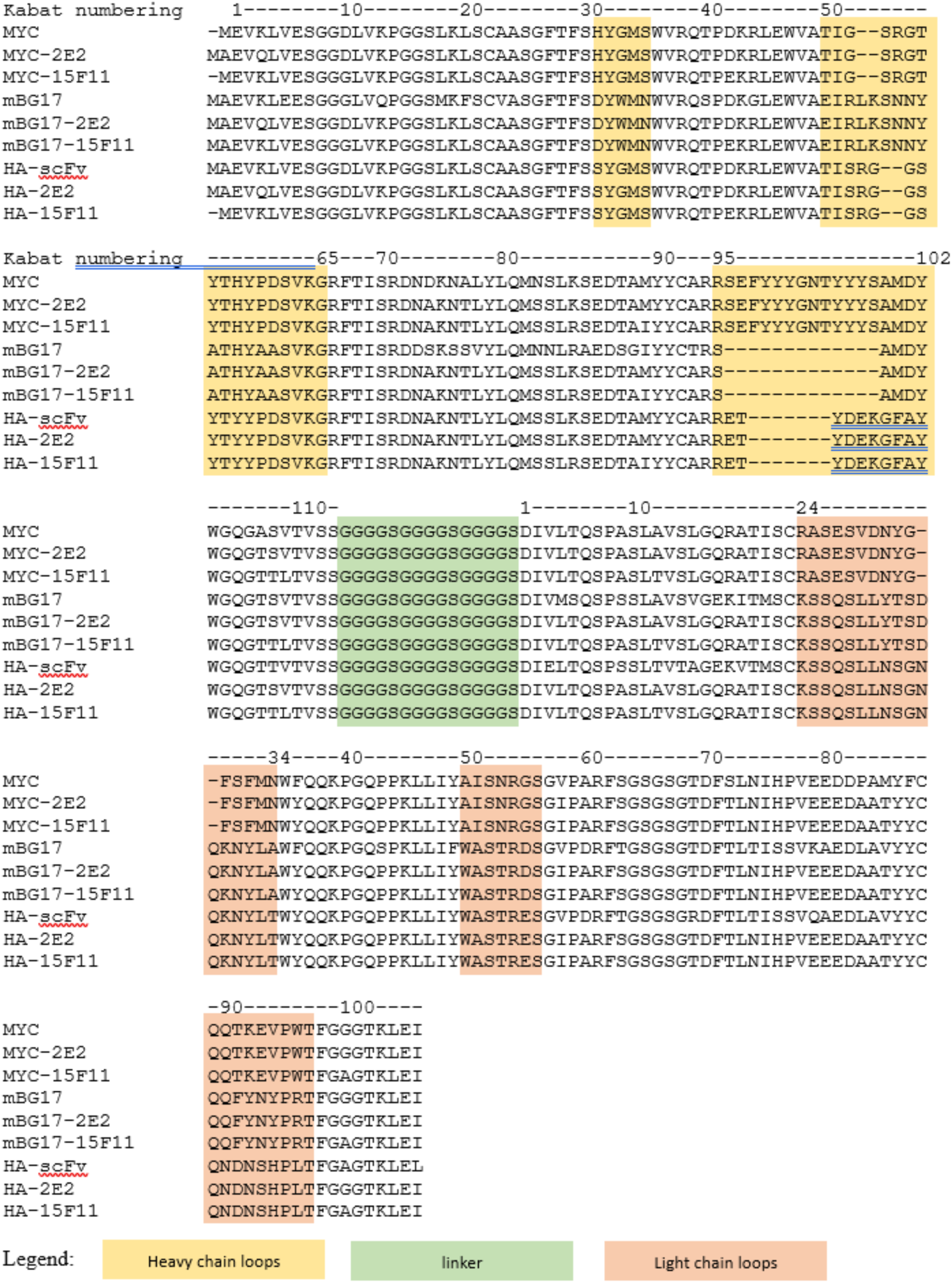

**Supplemental Figure 1.**
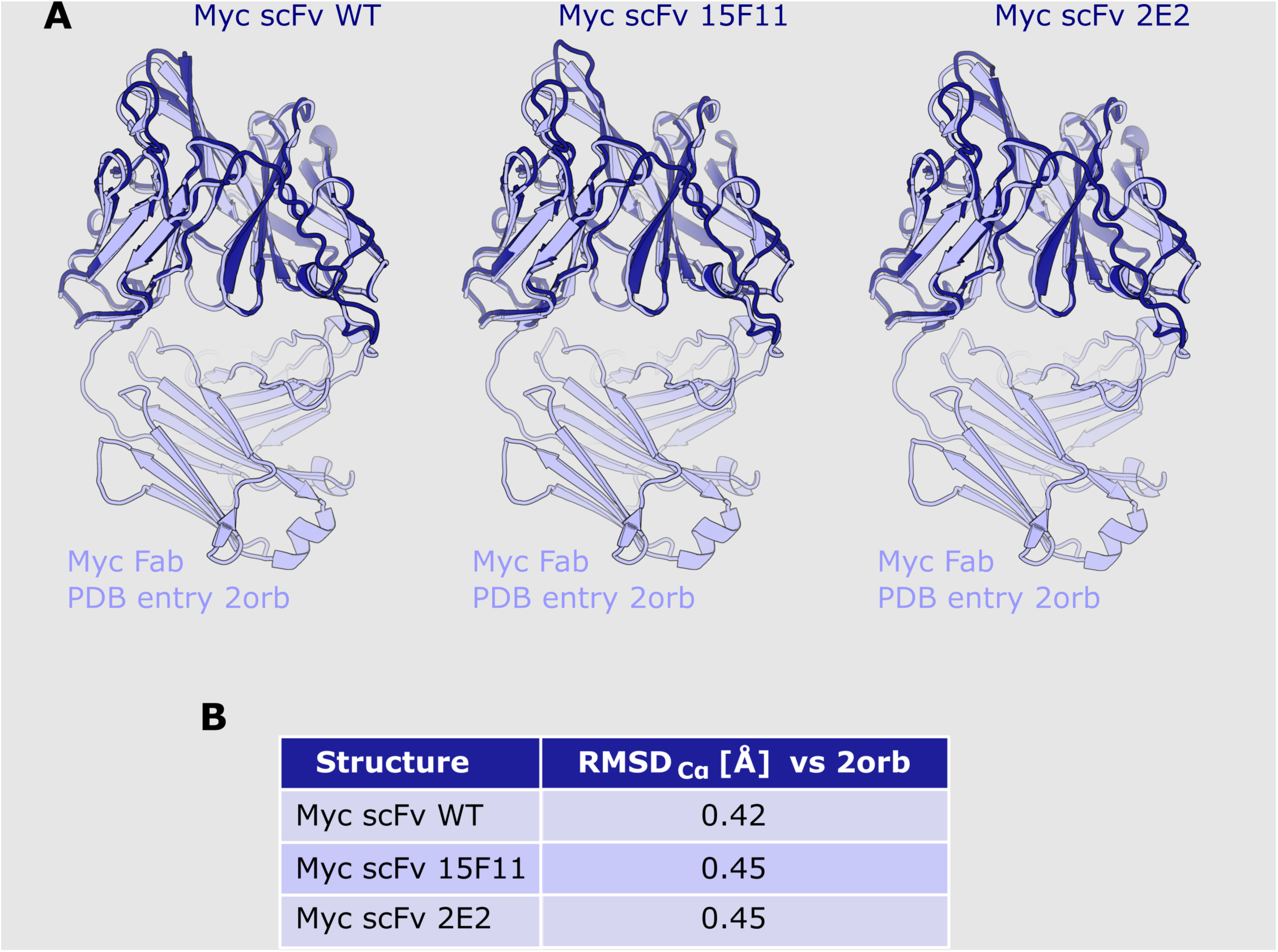
Alignment of AlphaFold2 predicted scFv structures to an anti-c-Myc Fab crystal structure. **A)** Alignments of AlphaFold2-derived wild-type Myc scFv, Myc-2E2 scFv, and Myc-15F11 scFv structures with a Myc Fab crystal structure (PDB: 2orb). Predicted scFv structures are shown in dark blue, 2orb Myc Fab structures are shown in light blue. **B)** RMSD values comparing structural similarities between the wild-type Myc scFv, Myc-2E2 scFv, and Myc-15F11 scFv structures with a Myc Fab crystal structure (PDB: 2orb) were computed by the PyMOL align command.

**Supplemental Figure 2:**
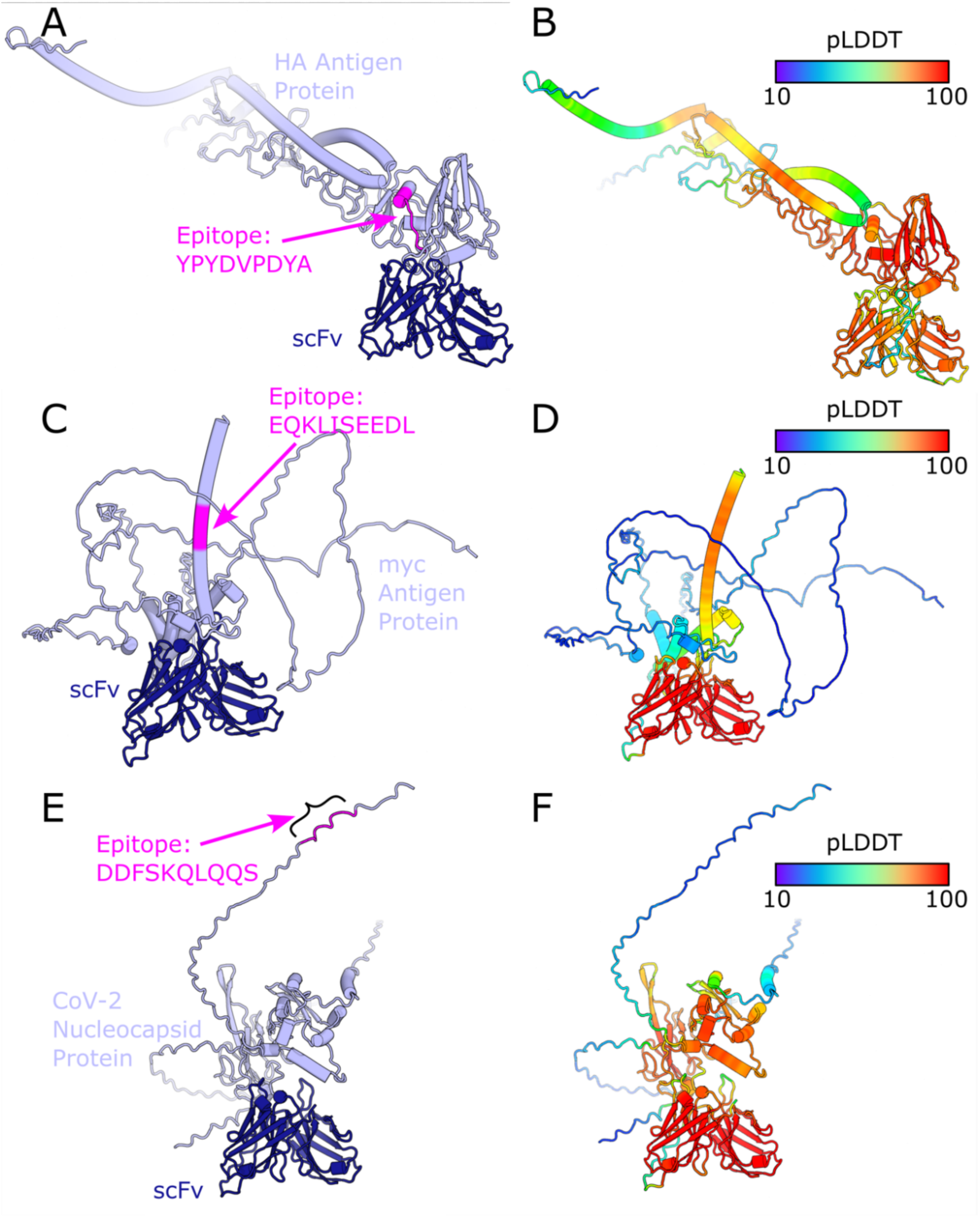
Alphafold2’s best attempt to dock whole sequences with the respective sequence’s scFv. **A)** The whole HA protein structure and scFv complex as predicted by AF2, with the correct epitope sequence highlighted in magenta. **B)** Shows the same structure by highlighted by confidence (pLDDT) of the structure with AF2. Similarly, the entire Myc protein-scFv complex are shown with **C)** the correct epitope highlighted in magenta and **D)** the confidence of the structure shown, and again for the mBG17 N-protein-scFv complex in **E)** and **F)**.

**Supplemental Figure 3:**
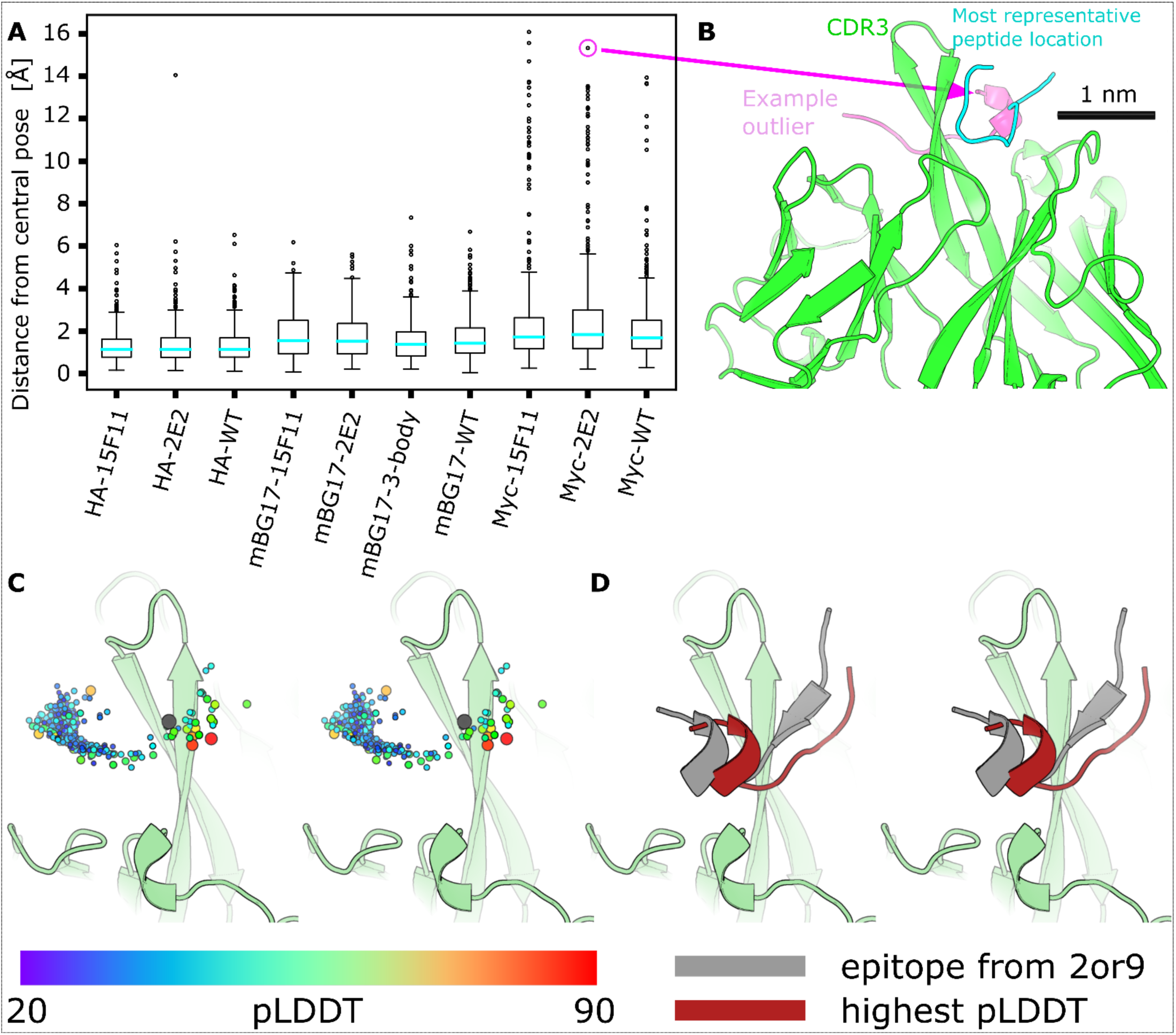
AlphaFold2 places all peptides near the CDR loops. The predicted Cα coordinates for all scFv (excluding the flexible linker) were extracted, and all were aligned together using the Kabsch algorithm (48, 49). With the scFvs structurally aligned, an all-against-all RMSD was calculated for the epitope peptides. To visually represent each peptide as a single point, the coordinates for all epitope atoms were averaged. The “central” exemplar epitope (cyan) is the peptide with the smallest sum of RMSD to all other peptides. **A)** The average and quartile for peptide placement relative to the central peptide via Box-and-Whisker plot reveals that AlphaFold2 largely places all epitopes in the same area. The Myc CDRH3 runs through the middle of a traditional paratope pocket, it isn’t a “cradle” for the epitope to sit on. AlphaFold2 places peptides on both sides of the CDRH3, causing significant spread in the peptide placement. **B)** An example of an exemplar, most-central predicted peptide structure (cyan) for the peptide PKSCASQDSS (cyan) bound to the Myc-2E2 scFv (green) that is distant from an example outlier peptide (magenta, peptide PHSPLVLKRC, center-to-center distance 14.8 Å). All peptide placements are still in contact with CDRH3, consistent with a strong AlphaFold2 bias to place peptides in a typical antibody binding site. **C)** The Myc-2E2 scFv (pale-green) and the average epitope placement (cyan) peptide alongside the crystal structure solution of the Myc epitope (grey). Remaining peptide placements are represented as a cloud of spheres at the mean peptide position. Each peptide sphere is colored and sized by epitope pLDDT (ranging from 20 to 90). Although AlphaFold2 frequently placed peptides on the opposite side of the CDRH3 from the Myc epitope (grey), it was not confident in these peptide placements (low, small, blue pLDDT spheres). In contrast, some of the peptides placed around the CDRH3, and in positions similar to the native epitope (grey) were placed with higher pLDDT confidence (increasingly large spheres trending from green to yellow to orange and red). **D)** The top ranked peptide as predicted by PAbFold with sequence QKLISEEDLL (red) and the crystal structure solution of the Myc epitope (grey).

**Supplemental Figure 4:**
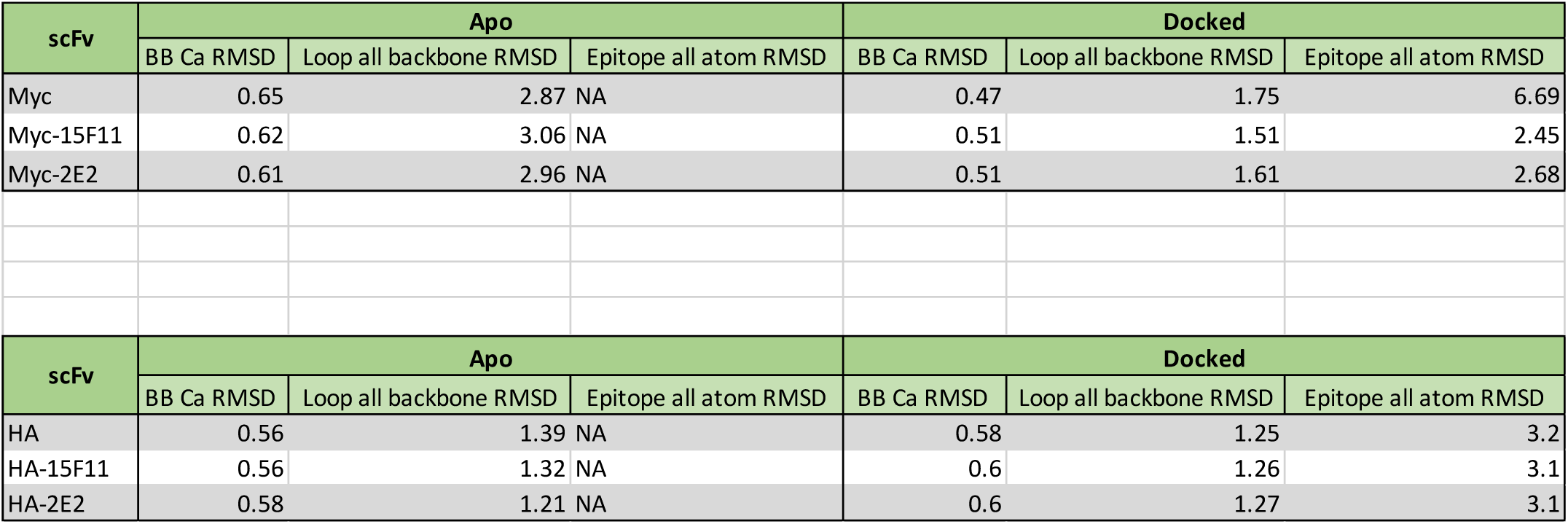
RMSD comparison (all numbers have units of Å) for AlphaFold2 predicted scFv structures compared to reference crystal structures, **A)** 2or9 (Myc) and **B)** 1frg (HA), respectively. The loops of the scFv more closely mimic the crystal structure when the epitope peptide is present. The backbone also undergoes subtle changes during docking that make it slightly more similar to the crystal structure. These structures were aligned by identifying the framework residues in all structures, then aligning the framework region Cα with the Kabsch algorithm (48, 49). Specifically excluded from this process were the heavy and light CDR loops of the structures, as well as the flexible linker structure that connects the heavy and light chains due to the inherent floppy, unstructured nature of this region. After aligning the framework regions of the AlphaFold2 predicted structures and the crystal structures (2or9 and 1frg respectively), an RMSD of these Cα was calculated and is reported as the first column ‘BB Cα RMSD’. Without further alignment, loop placement was analyzed with an all backbone RMSD by calculating the RMSD between the C, Cα, N, and O along the backbone of all residues in the scFv that were not used for the framework superimposition. This RMSD is reported in the second column as ‘Loop all backbone RMSD’. Finally, to investigate peptide predicted placement and potential scFv:epitope interactions, an all-atom RMSD was calculated between the crystal structure and the AF2 predicted peptide structure (no additional alignment). Because the apo structure lacks a peptide position, this is only reported in the ‘Docked’ category and is in the 3^rd^ column labeled ‘Epitope all atom RMSD’. One script was written for each scFv (Myc and HA), and can be found in the Zenodo deposition of our data (https://zenodo.org/records/10884181) because this analysis is not a key part of PAbFold. Briefly this analysis reveals that all three HA scFv variants have predicted framework regions and loop regions in the apo structures that closely match the reference structure (0.56-0.58 Å and 1.21-1.39 Å). Accordingly, when the cognate epitope peptide is present, it can be placed with relatively high accuracy for all three scFvs (3.1-3.2 Å), with only small changes in the loops (1.39 Å to 1.25 Å, 1.32 Å to 1.26 Å, and 1.21 Å to 1.27 Å). In contrast, the apo structures for the three Myc scFvs have a much higher deviation in the loop regions (2.87 to 3.06 Å). When the epitope peptide is added, there is significant motion in the loops consistent with an “induced fit” description. In the two chimeric Myc scFvs (Myc-15F11 and Myc-2E2) the final loop RMSD is reduced to 1.51-1.61 Å, and the epitope peptide is successfully predicted (2.45-2.68 Å). However, despite a lower apo-state loop RMSD (2.87 Å), the loop RMSD for the wild-type Myc scFv only drops to 1.75 Å, and the epitope peptide placement does not match the experimental structure (6.69 Å). This is consistent with the failure of the wild-type Myc scFv AlphaFold2 predictions in main text Figure 2.

**Supplemental Figure 5.**
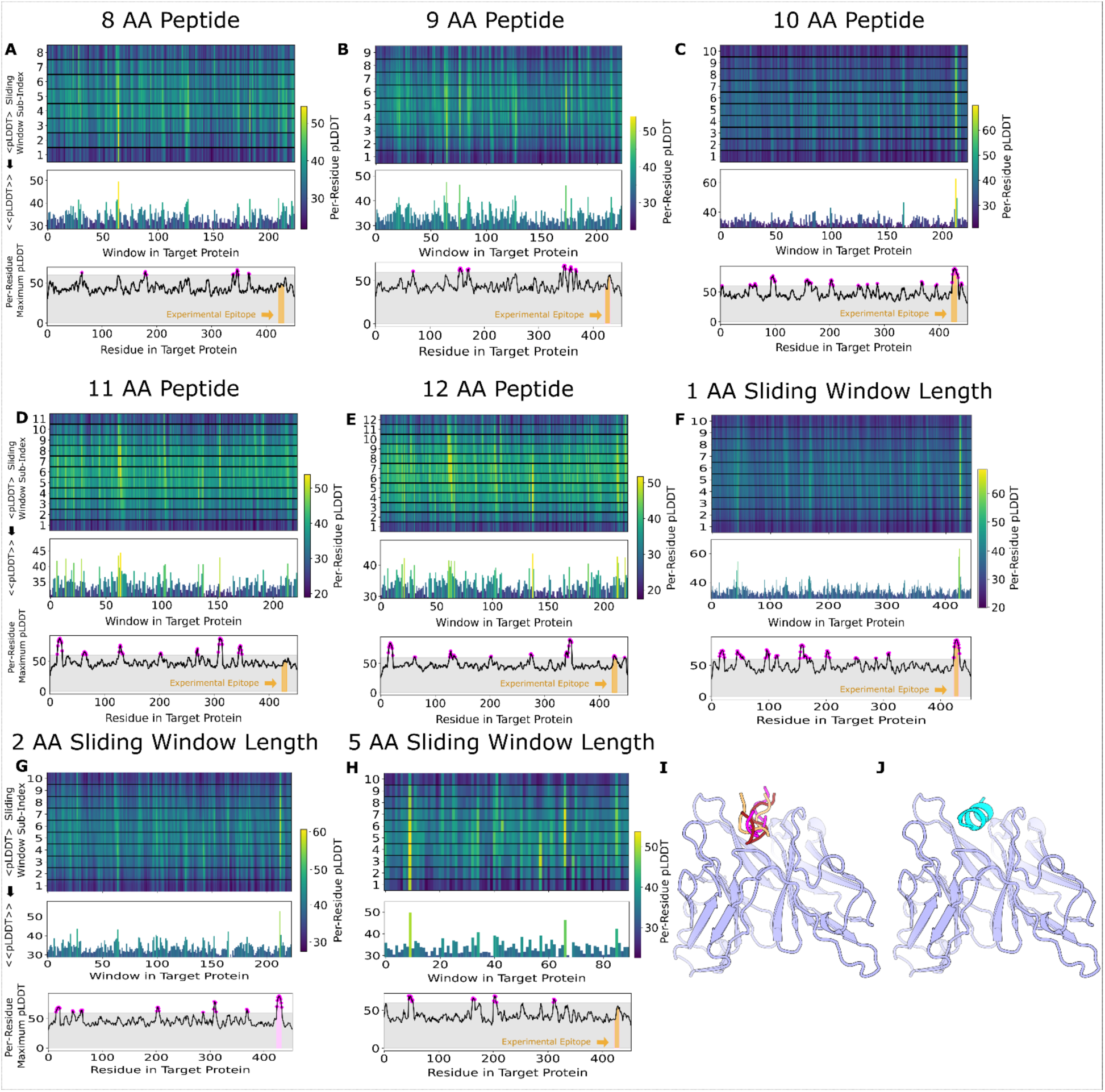
Assessment of peptide size and sliding window sizes on epitope prediction efficacy. Myc-2E2 scFv:peptide structures were predicted with peptides of 8 (**A**), 9 (**B**), 10 (**C**), 11 (**D**), and 12 (**E**) amino acid lengths derived from the Myc protein with a sliding window of 2 amino acids, and pLDDT scores from each predicted structure were plotted against the Myc amino acid position and sliding window length target. **F)** Negative control peptides bind to antibody binding sites, but with poor pLDDT scores. Similarly, with a fixed peptide length of 10 and a sliding window step size of 1 (**F**), 2 (**G**), and 5 (**H**), we can see the practical epitope detection outcome was similar for a sliding window of 1 and 2, but resolution and accuracy were reduced for a sliding window step size of 5. To more fully illustrate the strong learned bias that AlphaFold2 has for placing any peptides among the CDR loops, we predicted the structure of Myc-2E2 in complex with several control peptides. These negative control peptides bind to the generally expected antibody binding site, but with poor pLDDT. **I)** GSx5 in magenta (GSGSGSGSGS) had a score (mean peptide from Simple Max method pLDDT) of 29.5. (GGGGS)_2_ in orange (GGGGSGGGGS) had a score of 31.9. G_10_ in red (GGGGGGGGGG) had a score of 33. Lastly, **J)** A_10_ in cyan (AAAAAAAAAA) had a score of 41 and is the only negative control peptide to have an alpha-helical secondary structure (presumably due to the increased alpha helical propensity of alanine).

**Supplemental Figure 6:**
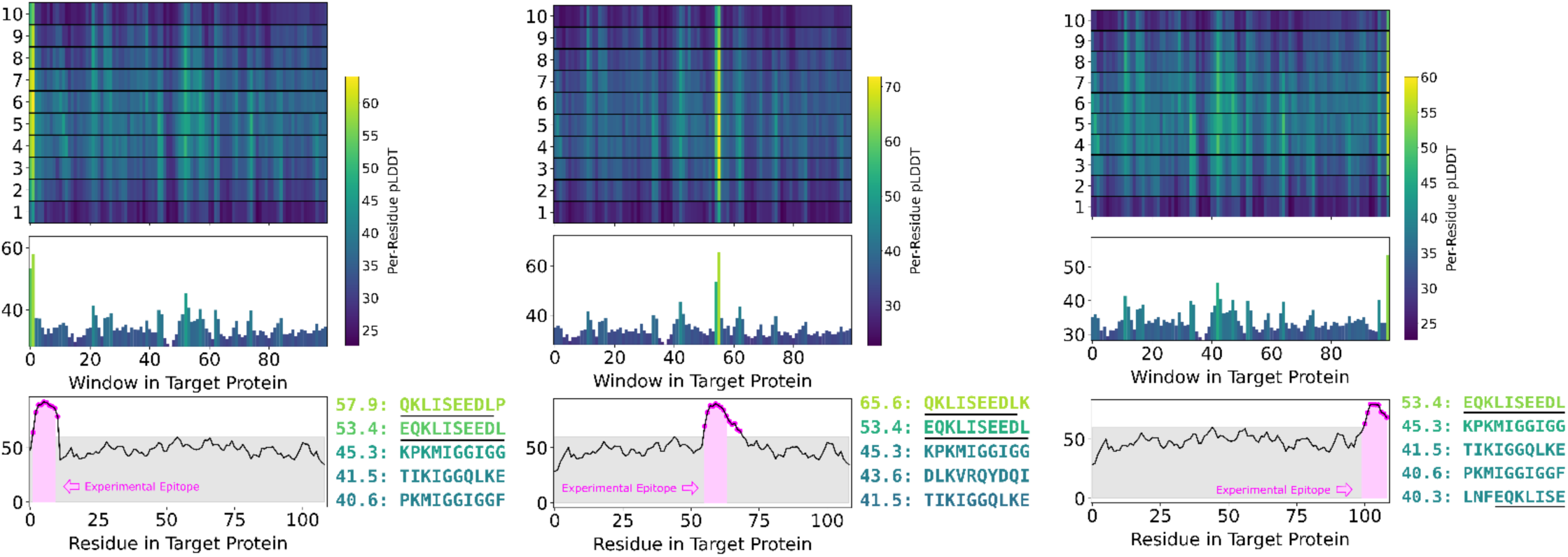
PAbFold epitope detection is independent of position within target sequence. The Myc epitope (EQKLISEEDL) was added into the beginning, middle, or end of the 99-a.a. HIV protease sequence (Genbank Accession: NP_705926.1) prior to epitope scanning structure prediction. Positions of the Myc epitope sequence added to in the **A)** N-terminus **B)** middle and **C)** C-terminus of the HIV protease sequence. **D)** Highlights the ranked sequences recovered from each experiment in A, B, and C.

**Supplemental Figure 7:**
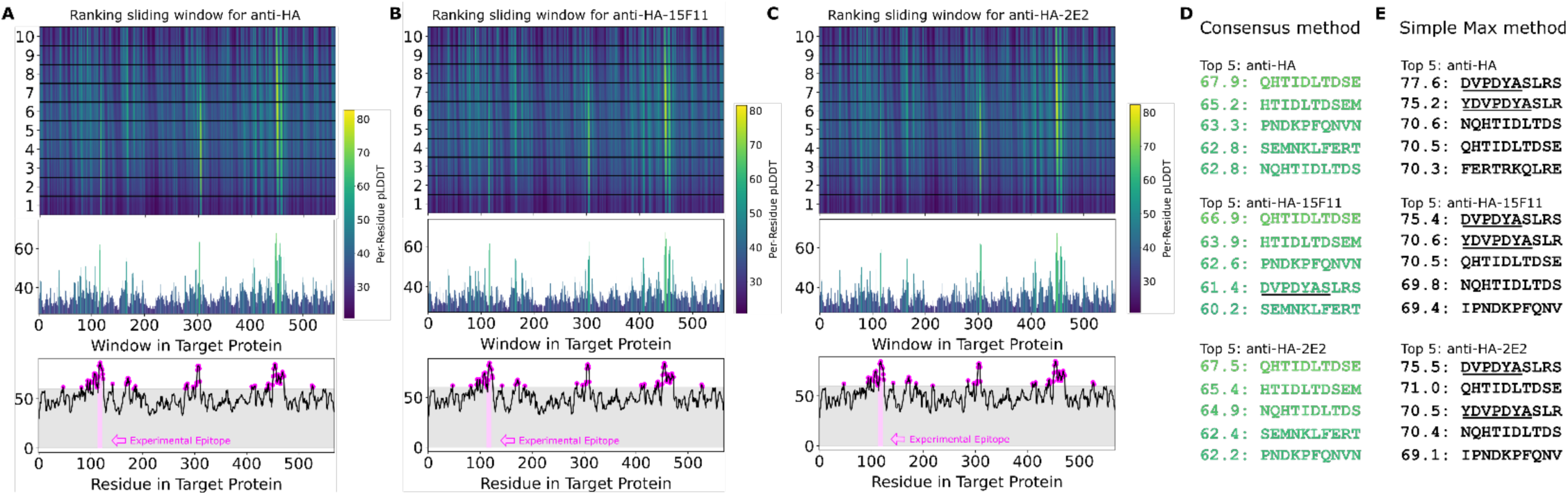
Alphafold2 can accurately predict the HA linear epitope in different scFv backbones. The anti-HA VH and VL antibody sequences were used to generate either **A)** wild-type scFv or CDR loop grafted onto the **B)** 15F11 or **C)** 2E2 antibody backbones. The Influenza A virus hemagglutinin protein sequence (Genbank AUT17530.1) was used as the target antigen and processed into 10 amino acid overlapping peptides with a 1 amino acid sliding window. The structures for each scFv:peptide pair were predicted with Alphafold2, and pLDDT values for each scFv:peptide pair are shown. **D)** The top-ranking epitope sequences via pLDDT scores are reported via the consensus method. Sequence underlining represents overlap with the known HA epitope (HA a.a. 114-125: YDVPDYASL). **E)** The top-ranking epitope sequences via pLDDT scores are reported via the simple max method.

**Supplemental Figure 8:**
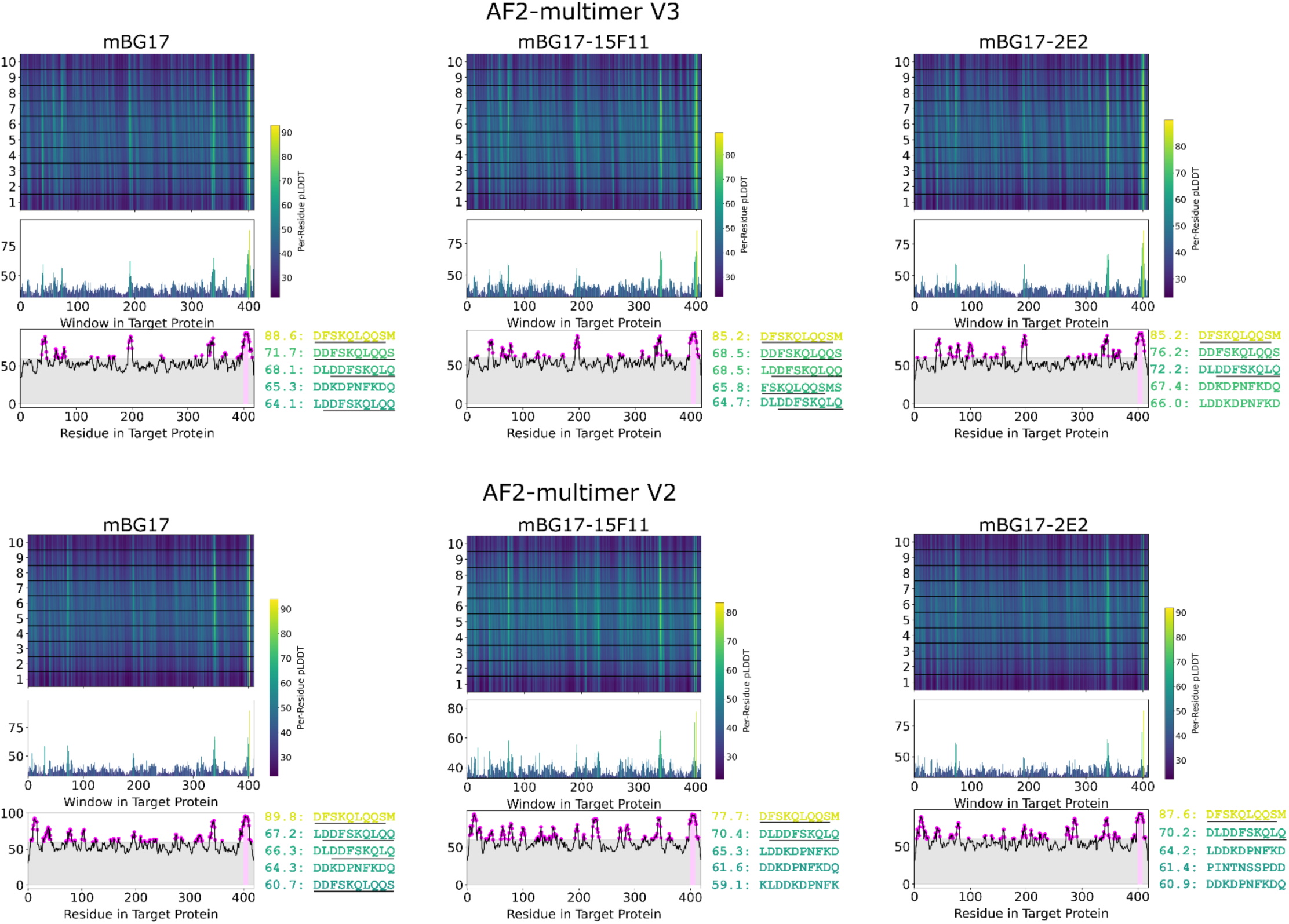
A comparison of Alphafold2 multimer version 3 and multimer version 2 applied to the mBG17 system. The experimental epitope, DDFSKQLQQS, is still easily identified with all three scFv backbones (wildtype, 15F11, and 2E2).

**Supplemental Figure 9:**
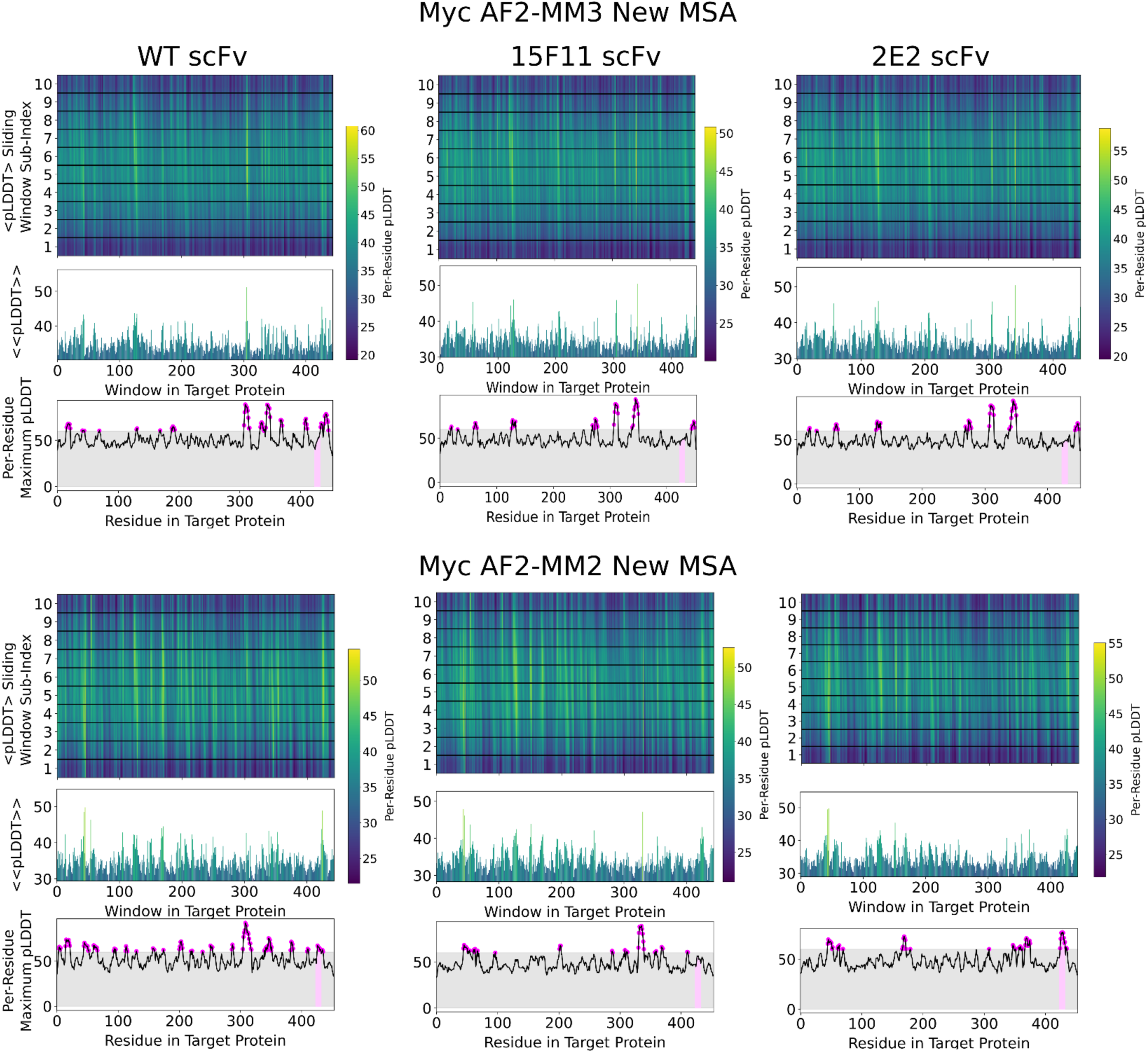
Myc comparison of epitope identification accuracy, comparing model types. Performance variation with AlphaFold2 model (multiple versions 2 and 3) and MSA versions (most up to date version of the ColabFold MSA server uses UniRef30 (2302) and PDB100 (220517)) vs the old MSA server (when this data was initially generated, ColabFold MSA server used UniRef30 (2202) and PDB70 (220313)). The left column is the WT scFv, the middle column is the CDR loops spliced onto the 15F11 backbone, and the right column is the CDR loops spliced onto the 2E2 backbone. Performance was ablated when using MM3 and the new MSA, and significantly degraded when using MM2 with the new MSA. For AF2-MM2 Old MSA, see Figure 2.

**Supplemental Figure 10:**
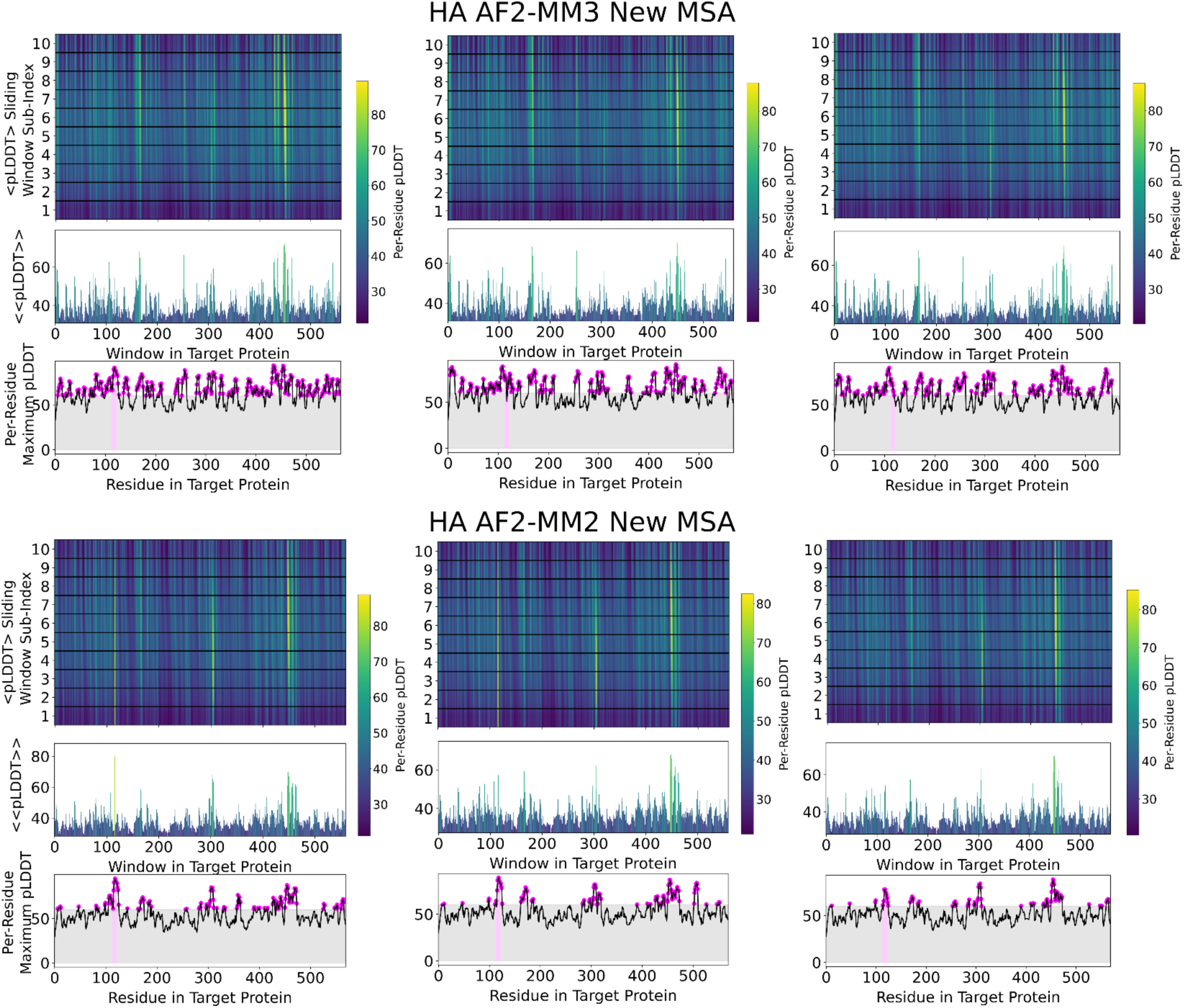
HA comparison of epitope identification accuracy, comparing model types. A comparison of the differing AlphaFold2 models with the Myc system (multimer version 3 and 2) along with a comparison of the new MSA (most up to date version of the ColabFold MSA server uses UniRef30 (2302) amd PDB100 (220517)) vs the old MSA server (when this data was initially generated, ColabFold MSA server used UniRef30 (2202) and PDB70 (220313)). The left column is the WT scFv, the middle column is the CDR loops spliced onto the 15F11 backbone, and the right column is the CDR loops spliced onto the 2E2 backbone. For AF2-MM2 Old MSA, see Supplemental Figure 7.

**Supplemental Figure 11:**
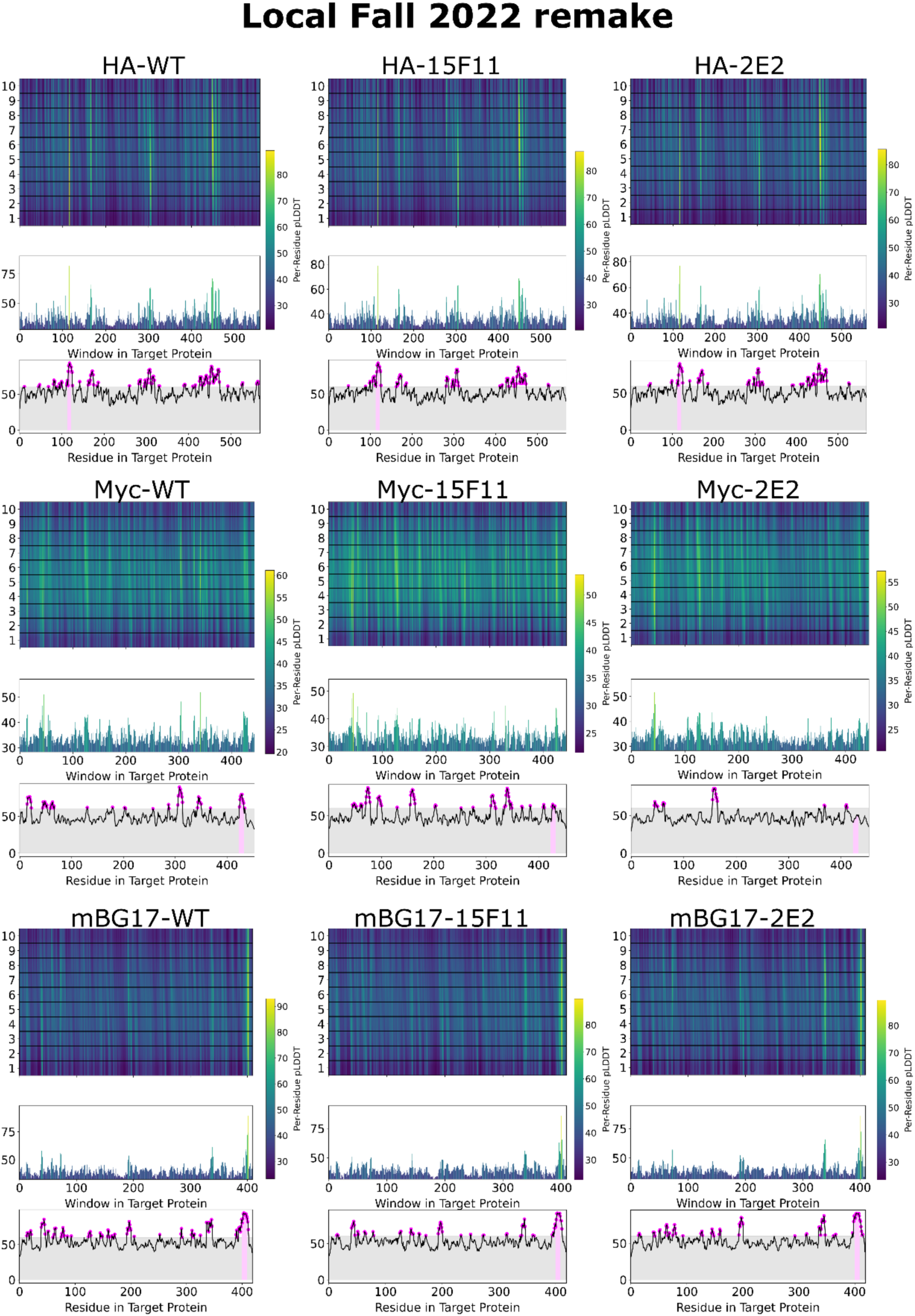
Local remake of the databases used by the MMSEQS server. Databases were downloaded (UniRef30 (2202) and PDB70 (220313)) and were queried locally to produced MSA’s for testing. These runs all were done with the multimer version 2 model of Alphafold 2. The left column is the WT scFv, the middle column is the CDR loops spliced onto the 15F11 backbone, and the right column is the CDR loops spliced onto the 2E2 backbone. The first row is the HA system, the second row is the Myc system, and the final row is the mBG17 system.

**Supplemental Figure 12:**
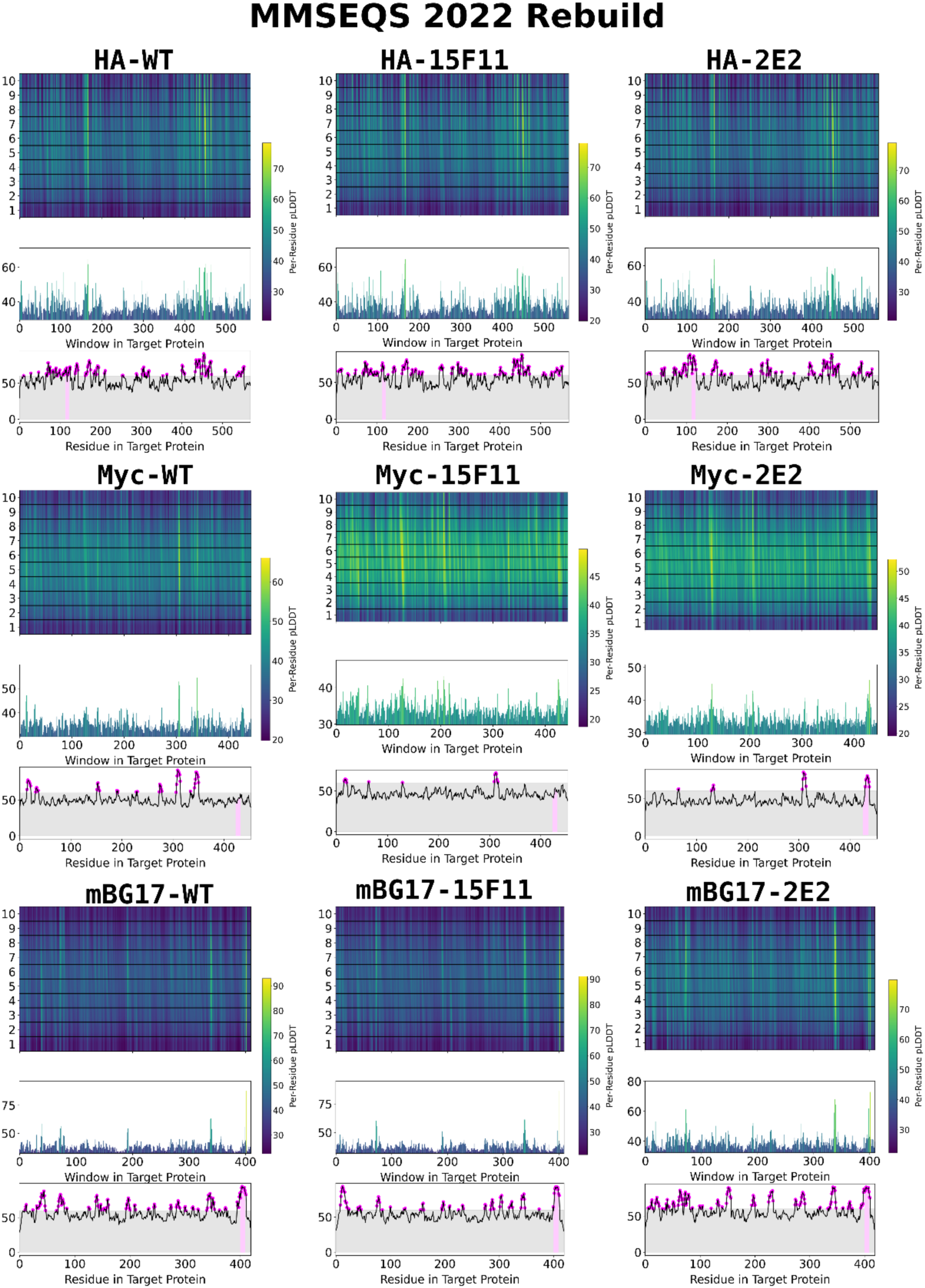
Server remake of the MMSEQS databases. The databases were rebuilt by the MMSEQS team UniRef30 (2202) and PDB70 (220313)) on the Colabfold MSA server and were queried produced MSA’s for testing. These runs all were done with the multimer version 2 model of Alphafold 2. The left column is the WT scFv, the middle column is the CDR loops spliced onto the 15F11 backbone, and the right column is the CDR loops spliced onto the 2E2 backbone. The first row is the HA system, the second row is the Myc system, and the final row is the mBG17 system.

**Supplemental Figure 13:**
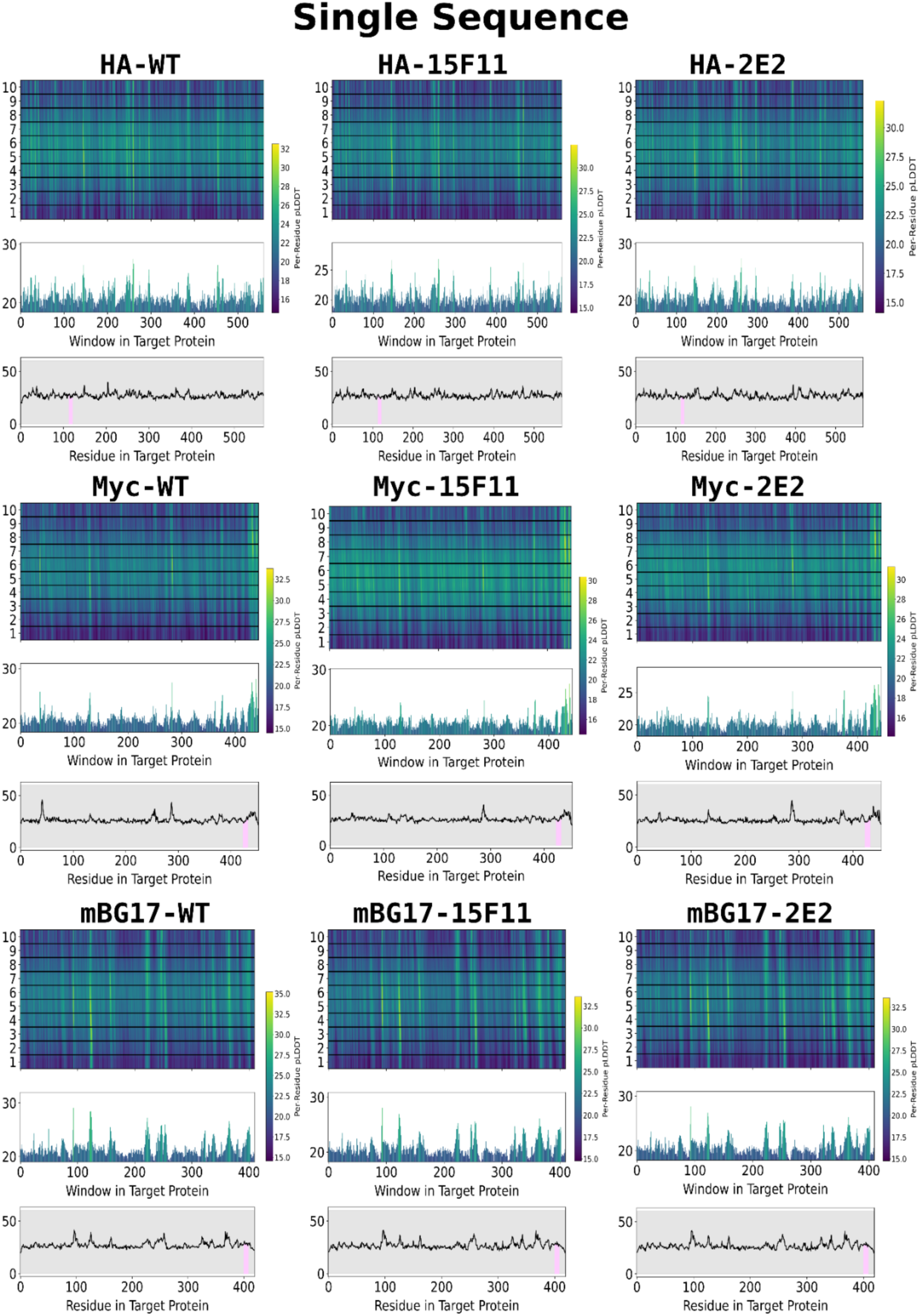
Single Sequence mode (no MSA’s) of epitope prediction with AF2. These runs all were done with the multimer version 2 model of Alphafold 2 in single sequence mode (i.e. no MSA was used) as a negative control, to highlight the importance of a quality MSA. The left column is the WT scFv, the middle column is the CDR loops spliced onto the 15F11 backbone, and the right column is the CDR loops spliced onto the 2E2 backbone. The first row is the HA system, the second row is the Myc system, and the final row is the mBG17 system.

**Supplemental Figure 14:**
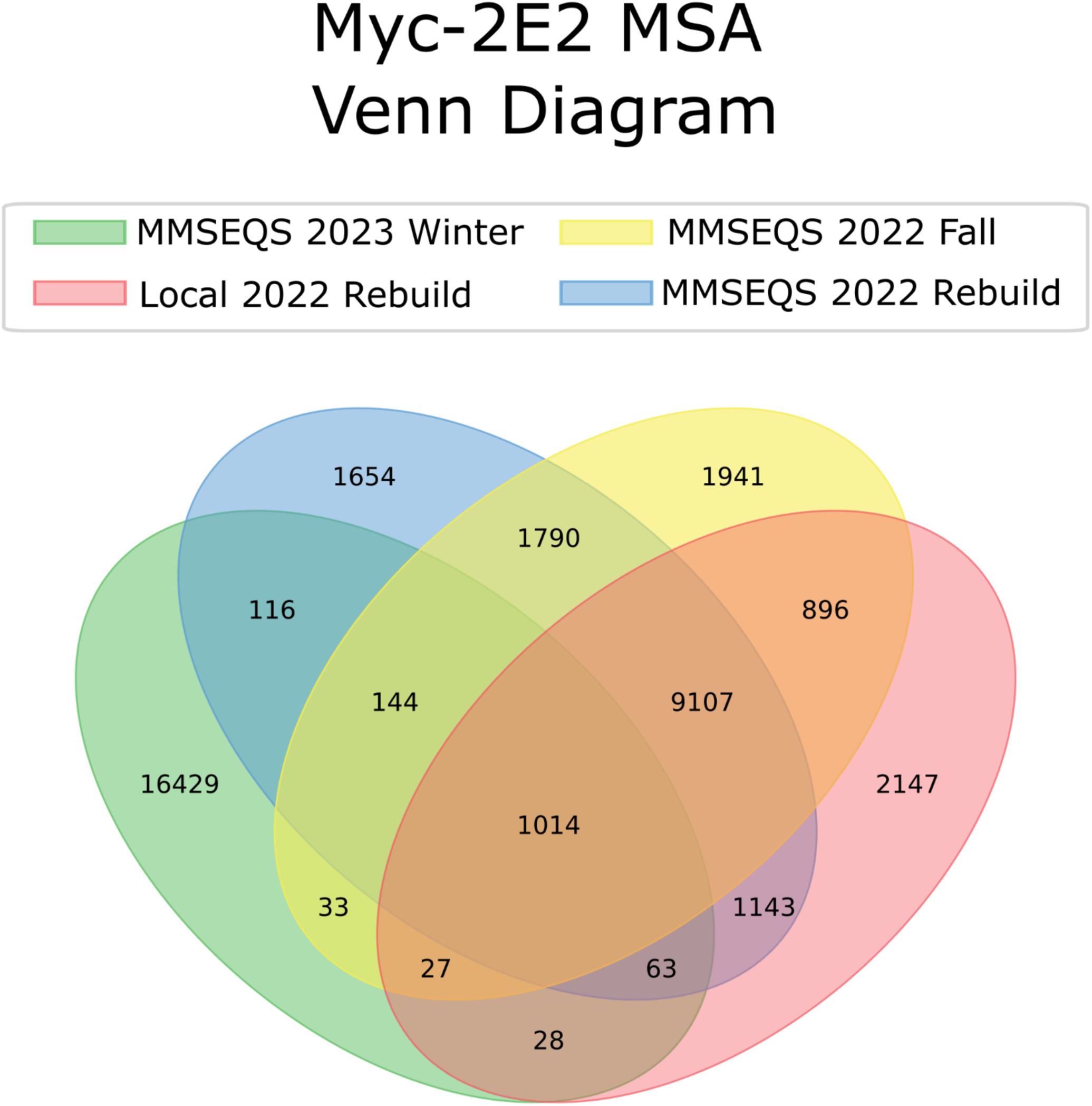
MSA overlap between the 4 generation methods. Here we highlight the number of unique entries that are shared amongst all of the MSA methods, those being: **1)** using the databases right now via colabfold (PDB30 2302 and PDB100 230517) (green) **2)** the databases after they had been accessed via colabfold and cached for repeated use (UniRef30 (2202) and PDB70 (220313)) (yellow), **3)** downloading the databases locally (UniRef30 (2202) and PDB70 (220313)) and attempting to create the MSAs ourselves (red), and **4)** querying the databases after the MMSEQS team rebuilt them for our use via colabfold (UniRef30 (2202) and PDB70 (220 313)) (blue).

**Supplemental Figure 15:**
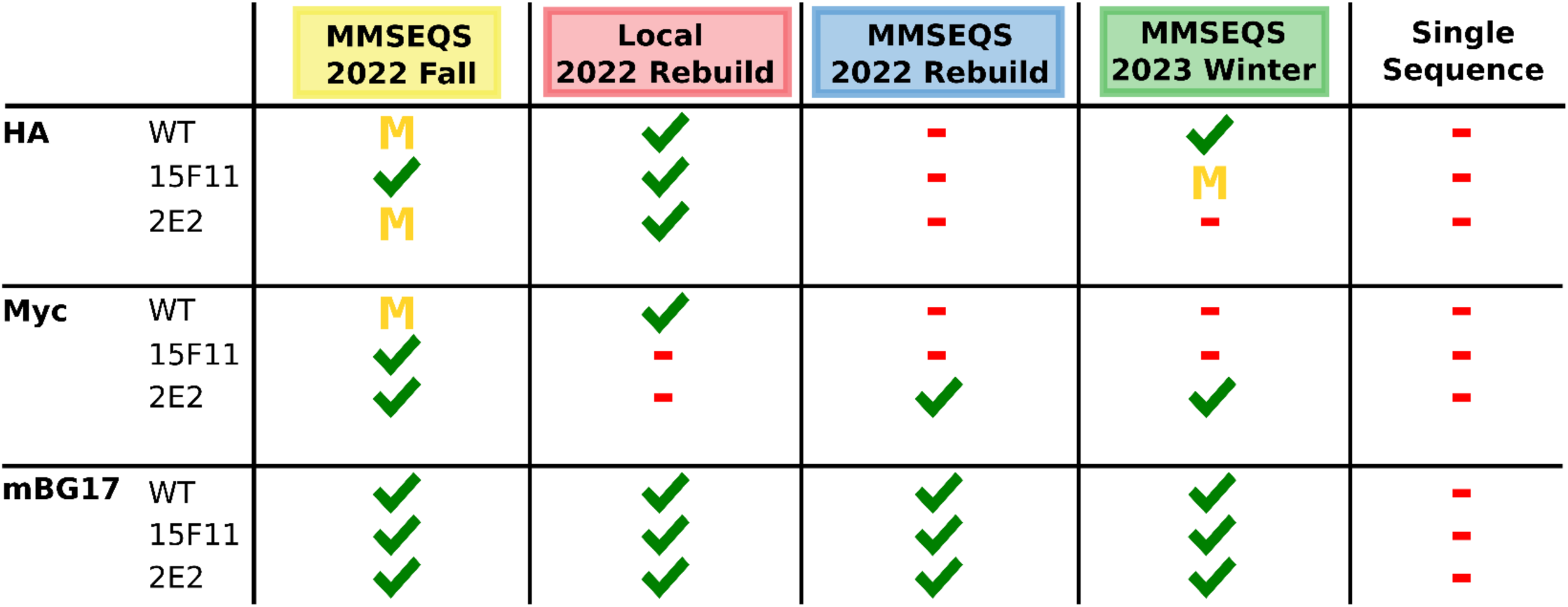
Comparison of how well each MSA generation scheme accurately identified the experimentally derived epitope within the top 5 epitope sequences. A green checkmark shows that it was found by both the consensus model and the top single model, a yellow “M” means the simple max method correctly identified the experimental epitope in the top 5 epitopes, and the red dash means both methods failed. The consensus model did not identify the epitope correctly when the simple max method failed to. The colored background behind the titles is the same color as Supplemental Figure 14 to help guide the eye.

